# Restoring multiple TDP-43 cryptic targets, but not solely *Unc13a*, rescues motor neuron disease

**DOI:** 10.64898/2026.09.11.751082

**Authors:** Aswathy Peethambaran Mallika, Meghraj Singh Baghel, Opal Sitzman, Jessica G. Yu, Tianyu Cao, Shruti Renganathan, Irika R. Sinha, Tatiana Melnikova, Jonathan P. Ling, Philip C. Wong

## Abstract

Dysfunction of TAR DNA-binding protein 43kDa (TDP-43) underlies amyotrophic lateral sclerosis (ALS), a neurodegenerative disorder with limited therapeutic options. While current therapeutic approaches are designed to individually target unique cryptic exons of TDP-43 such as *UNC13A*, the sufficiency of such a strategy to mitigate motor neuron disease remains unclear. Using a mouse model lacking TDP-43 in spinal motor neurons which mimics early stages of ALS, we show that the exclusion of *Unc13a* cryptic exon fails to mitigate motor neuron disease. In contrast, the restoration of multiple TDP-43 cryptic targets, including *Unc13a*, attenuated motor neuron loss, and rescued motor neuron disease. Additionally, compared to brain neurons, spinal motor neurons accumulate markedly lower amounts of *Unc13a* cryptic exons in mice and humans, suggesting that the contribution of this TDP-43 cryptic target to spinal motor neuron loss may be limited. Together, these results strongly support ALS therapeutic strategies designed to simultaneously restore multiple TDP-43 cryptic targets to attenuate spinal motor neuron loss.

---

Emerging evidence indicates that deficits associated with TAR DNA-binding protein 43 (TDP-43) cryptic splicing^1–7^ are a central pathogenic mechanism underlying Amyotrophic Lateral Sclerosis-Frontotemporal Dementia (ALS-FTD). TDP-43, an RNA-binding protein, was first linked to ALS-FTD through the observation of cytoplasmic aggregates in end-stage disease^8^. A key function of TDP-43 is to repress cryptic exons, intronic sequences whose inclusion disrupts maturation of mRNA transcriptome-wide which is thought to contribute to neuron loss^7^. Deficits in TDP-43 splicing repression can be detected during presymptomatic stage in biofluids of ALS-FTD patients^1^, and such dysfunction is thought to precede its cytoplasmic aggregates by at least a decade in the aging human brain^9^, suggesting that loss of TDP-43 splicing repression drives neuron loss^9^. This interpretation is strengthened by evidence that ALS-linked TDP-43 mutations^10–12^ promote the inclusion of the *STMN2* cryptic exon in human iPSC-derived neurons, independent of TDP-43 cytoplasmic aggregates^5,6,13^. A major risk allele for ALS-FTD that promotes the inclusion of a cryptic exon in *UNC13A* normally repressed by TDP-43 could also account for earlier disease onset and progression of ALS-FTD^3,4,14^, suggesting the possibility of preventing cryptic splicing of these targets relevant for neuronal function may slow disease. While therapeutic strategies designed to prevent cryptic splicing of either *STMN2* or *UNC13A* using the antisense oligonucleotide (ASO) platform have recently entered clinical evaluation, the sufficiency of restoring cryptic splicing of either target alone within the context of TDP-43-associated neurodegenerative dysfunction has not been documented. In the case of *UNC13A in vitro* studies designed to evaluate the sufficiency of preventing its cryptic splicing, remain inconclusive^15–17^. Clarification on the efficiency of preventing multiple cryptic targets and/or solely *UNC13A* in attenuating motor neuron disease will impact development of therapeutic strategies designed to slow the progression of ALS.

To address this question, we took a two-pronged genetic approach. First, we deleted a nonconserved TDP-43-regulated cryptic exon in *Unc13a* (*Unc13aCE^-/-^*mice). Although this cryptic exon is located at a different location than the one in human *UNC13A*, it is predicted to similarly reduce the expression of mouse *Unc13a*^18^ as occurs in *UNC13A* due to the inclusion of the human-specific cryptic exon^3,4^. By selectively ablating *Tardbp* in spinal motor neurons (*ChAT-Cre;Tardbp^f/f^*, referred here as TDP-43cKO mice)^7^ of *Unc13aCE^-/-^* mice, we tested whether preventing cryptic splicing of *Unc13a* alone is sufficient to attenuate motor neuron disease. Second, to determine the sufficiency of restoring multiple TDP-43 targets, including *Unc13a*, to rescue motor neuron disease, we used a BBB-crossing AAV serotype (AAV-PhP.eB) capable of robustly delivering a previously established splicing repressor (CTR)^19^.

## RESULTS

### Prevention of *Unc13a* cryptic splicing fails to attenuate motor neuron disease

To test whether the prevention of cryptic splicing of *Unc13a* alone is sufficient to mitigate motor neuron disease, we used a CRISPR-Cas 9 approach to delete a 44bp fragment from the mouse genome that defined the *Unc13a* cryptic exon (**Fig. 1a, Ext Data Fig. 1e**). To confirm the deletion, we used PCR analysis and RNA *in situ* hybridization (BaseScope-ISH RED assay) to detect the presence of the *Unc13a* cryptic exon (**Fig. 1b**). As a result of this deletion, *Unc13a* transcripts are incapable of encoding the cryptic exon, regardless of TDP-43 expression levels, and transcribed mRNA does not include the noncanonical cryptic sequence. To determine the impact of preventing *Unc13a* cryptic exon inclusion within the context of TDP-43 depletion and the motor neuron disease phenotype, *Unc13aCE^-/-^*mice were crossbred with *ChAT-Cre;Tardbp^f/f^* mice to generate a cohort of mice with motor neuron-specific TDP-43 depletion but lacking the *Unc13a* cryptic exon inclusion (*ChAT-Cre;Tardbp^f/f^*;*Unc13aCE^-/-^*). Grip strength and body weight analysis of *ChAT-Cre;Tardbp^f/f^*;*Unc13aCE^-/-^*mice along with littermate *ChAT-Cre;Tardbp^f/f^* mice revealed *ChAT-Cre;Tardbp^f/f^*;*Unc13aCE^-/-^* mice could not maintain the normal grip strength and body weight as compared to those of the age matched controls (**Fig.1 e, f**), which progressed to hind limb paralysis between 6-8 months of age and became moribund between 6-9 months of age, phenotypes that are indistinguishable from those observed in *ChAT-Cre;Tardbp^f/f^* mice (**Fig. 1g**). Pathological analysis of motor neurons of 5-month-old *ChAT-Cre;Tardbp^f/f^*;*Unc13aCE^-/-^*mice revealed significant loss of motor neurons like that found in *ChAT-Cre;Tardbp^f/f^*mice (**Fig. 1c, d**). Taken together, these results suggest that preventing cryptic splicing of *Unc13a* is insufficient to attenuate motor neuron disease.

**Fig. 1:**
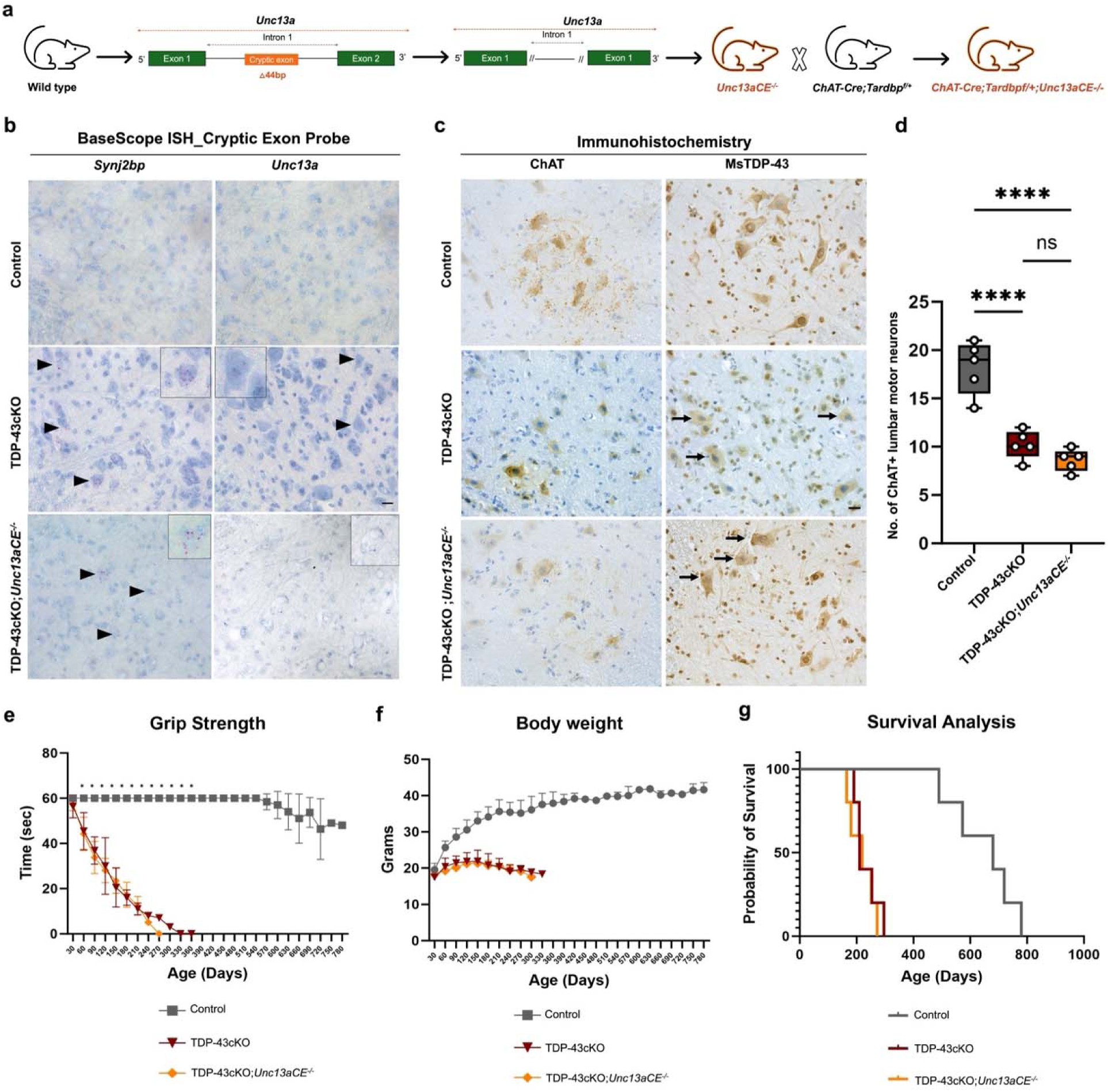
Deletion of *Unc13a* cryptic exon fails to rescue the motor neuron disease phenotype in *ChAT-Cre;Tardbp^f/f^* (TDP-43cKO) mice. (**a**) Schematic showing the creation of mouse model with *Unc13a* cryptic exon within intron 1 (**b**) Representative images of BaseScope-ISH RED assay performed on lumbar spinal cord sections targeting the cryptic exons in *Synj2bp* and *Unc13*a in control, TDP-43cKO and TDP-43cKO;*Unc13aCE^-/-^* mice. Arrow heads point to cells with cryptic exons. *Unc13a* cryptic exon expression was low in the TDP-43 depleted motor neurons as compared to the expression of *Synj2bp* in TDP-43cKO mice (n=3, 1m old). Scale bar=20 µm. (**c,d**) Pathologic analysis of 5m old mice (n=5 per group). Immunohistochemical analysis using antibodies against endogenous TDP-43 and ChAT in lumbar motor neurons lacking TDP-43. Scale bar=20 µm. (**d**) Quantification of number of surviving lumbar motor neurons in 5m old control, TDP-43cKO and TDP-43cKO;*Unc13aCE^-/-^* mice (n=5 per group). (**e-g**) Phenotypic analysis including grip strength (**e**), body weight (**f**), and survival analysis (**g**). n=5 per group.

### Intravenous delivery of AAV-PHP.eB-CTR to symptomatic *ChAT-Cre;Tardbp^f^*^/f^ mice efficiently transduces motor neurons

To determine whether restoring multiple cryptic targets of TDP-43 attenuates neuron loss and rescues motor neuron disease, we utilized our *ChAT-Cre;Tardbp^f/f^*mouse line, in which the *ChAT-Cre* driver selectively deletes TDP-43 in motor neurons^19^. These mice develop a progressive motor neuron disease that ultimately leads to premature death. Using a previously established breeding scheme, we generated a large cohort of *ChAT-Cre;Tardbp^f/f^*mice and littermate controls (see materials and methods & **Extended Data Fig.1**). CTR-a <u>C</u>himeric <u>T</u>DP-43 splicing <u>R</u>epressor designed by combining the TDP-43 RNA-recognition domain with another splicing regulator, *RAVER1* (replacing the aggregation prone low-complexity domain in TDP-43, **Extended Data Fig.1**)^7,19^-was delivered by lateral tail vein injection of AAV-PHP.eB-CTR at the optimized dose of 5E+13 vg/Kg (**Extended Data Fig. 2**). Injections were performed at 6 weeks of age, when fine tremors are already apparent in *ChAT-Cre;Tardbp^f/f^*mice, resulting in accumulation of CTR in motor neurons throughout the brain stem and spinal cord. The transduction rate of motor neurons in lumbar spinal cord and brain stem was 79±6.2% and 72±11.7%, respectively, as determined by CTR immunostaining and quantification at one-month post-injection (**Fig. 2b & Extended Data Fig. 3, 4**).

**Fig. 2:**
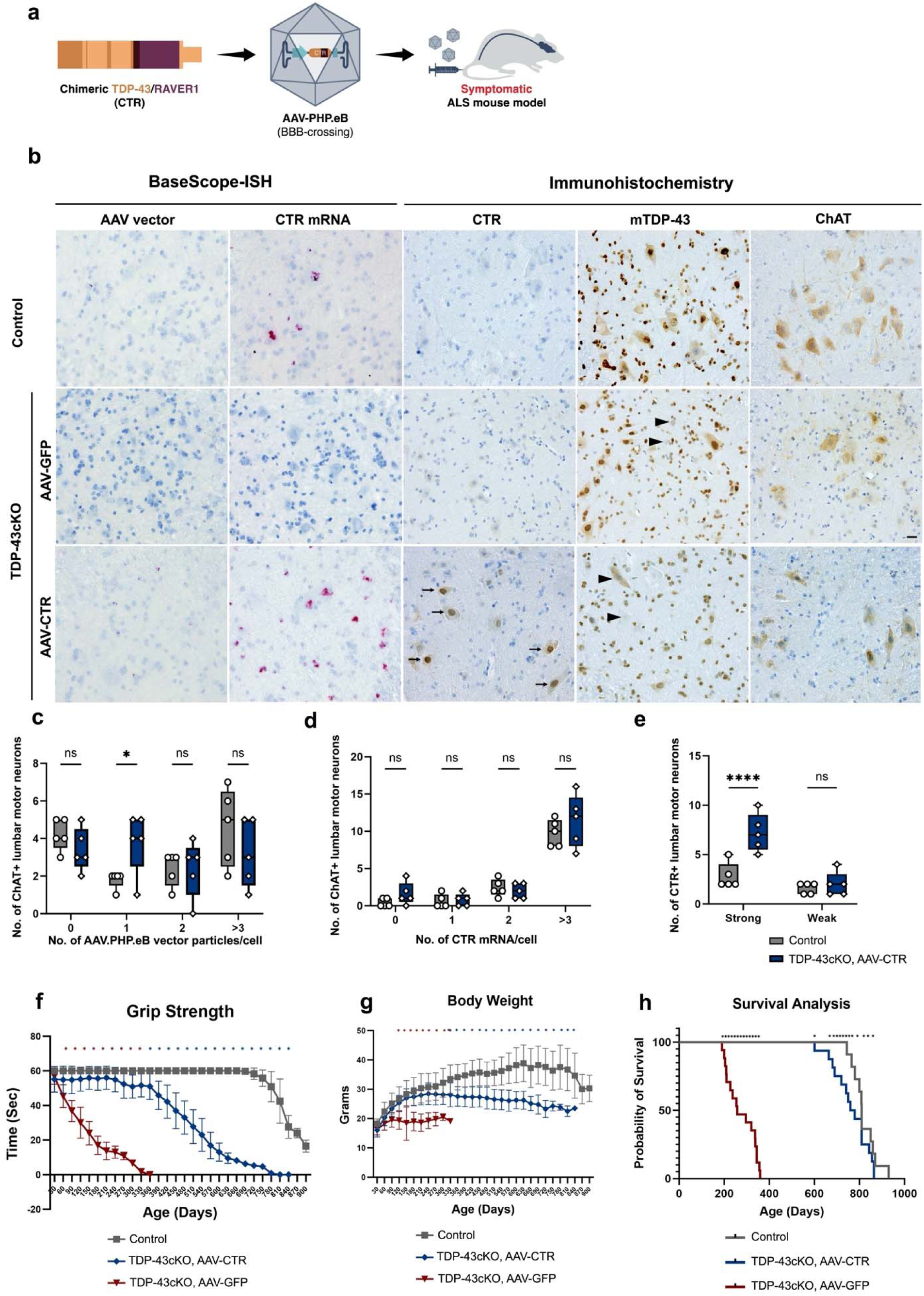
Intravenous delivery of AAV-PHP.eB-CTR confers high efficiency of transduction in adult motor neurons. (**a**) Schematic diagram showing the study design. (**b**) Representative images showing AAV.PHP.eB viral vector particles (red dots), CTR mRNA particles (red dots) and the accumulation of CTR protein in ventral motor neurons of TDP-43cKO (*ChAT-Cre;Tardbp^f/f^*) mice. Immunostaining for CTR was carried out using the anti-human TDP-43 antibody in lumbar spinal cord sections of 3-month-old TDP-43cKO mice (indicated by arrows in Panel 3). Intravenous administration of AAV-PHP.eB-CTR results in a mean infectivity rate of 79±6.2% (n=5). Immunostaining for mouse endogenous TDP-43 (mTDP-43) indicates ∼60% depletion of endogenous TDP-43 (indicated by arrowheads in Panels 2 and 3). Panel 1: AAV-PHP.eB-CTR treated control (*ChAT-Cre;Tardbp^f/+^*) mice; panel 2: AAV-PHP.eB-GFP treated TDP-43cKO (*ChAT-Cre;Tardbp^f/f^*) mice; and panel 3: AAV-PHP.eB-CTR treated TDP-43cKO (*ChAT-Cre;Tardbp^f/f^*) mice. n=5 per group. Scale bar=20 µm. Single cell quantification of AAV genome copy number (**c**) and CTR mRNA expression (**d**) using BaseScope-ISH probes and immunohistochemical staining of CTR (**e**) shows that the autoregulatory domain expressed along with CTR prevents its overexpression in cells with normal expression of TDP-43. n=5 per group. (**f,g,h**) Efficacy of intravenous delivery of AAV-PHP.eB-CTR to TDP-43cKO mice. (**f**) Grip strength analysis revealed marked amelioration in motor deficits (characteristic of the TDP-43cKO mice) in AAV-PHP.eB-CTR treated adult TDP-43cKO mice (*p<0.0001, two-way ANOVA test). (**g**) Intravenous administration of AAV-PHP.eB-CTR shows significant improvement in the age-dependent reduction of body weight associated with the TDP-43cKO mice (*p<0.0001, two-way ANOVA using Tukey’s multiple comparison test). (**h**) Intravenous administration of AAV-PHP.eB-CTR extended survival in TDP-43cKO mice. Kaplan-Meier survival curves are shown for control, AAV-PHP.eB-CTR and AAV-PHP.eB-GFP treated TDP-43cKO mice. Data from male and female cohorts aged between 23-28 months are shown together. n=15 per group.

### Inclusion of TDP-43 autoregulatory domain regulates CTR protein levels and prevents overexpression toxicity

CTR functionally mimics the splicing repression activity of TDP-43 and reduces aberrant splicing events at target genes. To prevent the potential toxicity of excessive accumulation of CTR, which would likely mimic the deleterious effects of TDP-43 overexpression^20–22^, we engineered AAV-PHP.eB-CTR to incorporate the endogenous autoregulatory element which is naturally present within the TDP-43 3’UTR as an intrinsic “safety switch”. RNA *in situ* hybridization analysis using specific probes revealed the presence of both the AAV backbone and CTR mRNA within the motor neurons of spinal cord and brainstem, which allowed determination of the number of AAV particles and CTR transcripts expressed within each tissue (**Fig. 2b, c, d & Extended Data Fig. 4**). While the genome copy number of AAV-PHP.eB-CTR is maintained at 2-3 copies per cell (**Fig. 2c**), the accumulation of CTR mRNA in spinal motor neurons was higher in the TDP-43 deficient motor neurons of *ChAT-Cre;Tardbp^f/f^* mice when compared to normal motor neurons in *ChAT-Cre;Tardbp^f/+^* mice (**Fig. 2d, Extended Data Fig. 4)**. These data indicate that CTR transcripts are suppressed when TDP-43 is present within a motor neuron and is thus successfully regulated by TDP-43 expression. Furthermore, CTR protein expression was elevated in spinal motor neurons lacking TDP-43 as compared to those in *ChAT-Cre;Tardbp^f/+^* mice (**Fig. 2e**). Long-term monitoring of control *ChAT-Cre;Tardbp^f/+^* mice treated with AAV-PHP.eB-CTR showed normal grip strength, body weight, and histopathology, with no evidence of vector-related toxicity. Together, these findings confirm that the TDP-43 3’UTR autoregulatory element can serve as an effective safety mechanism to prevent over-accumulation of CTR protein in motor neurons.

### Delivery of AAV-PHP.eB-CTR to symptomatic *ChAT-Cre;Tardbp^f/f^* mice rescues motor neuron disease

To determine the impact of AAV-PHP.eB-CTR on motor neuron disease phenotype, we initially assessed the grip strength of *ChAT-Cre;Tardbp^f/f^*mice treated at a symptomatic stage. As expected, AAV-PHP.eB-GFP treated *ChAT-Cre;Tardbp^f/f^*mice showed deficits as early as 6 weeks of age and continued to worsen over the next 6 months (**Fig. 2f**). Remarkably, *ChAT-Cre;Tardbp^f/f^*mice treated with AAV-PHP.eB-CTR maintained normal grip strength over this same period of monitoring as compared to control *ChAT-Cre;Tardbp^f/+^*mice (**Fig. 2f**). In addition to this rescue of grip strength, the body weight of AAV-PHP.eB-CTR treated *ChAT-Cre;Tardbp^f/f^* mice also continued to increase, like that of control littermates (**Fig. 2g & Extended Data Fig. 5**). In contrast, the body weight of AAV-PHP.eB-GFP injected *ChAT-Cre;Tardbp^f/f^* mice failed to increase over this same period (**Fig. 2g & Extended Data Fig. 5**). To determine whether improved grip strength and the ability to maintain normal body weight impacted survival of AAV-PHP.eB-CTR treated *ChAT-Cre;Tardbp^f/f^*mice, we monitored the progression of disease until AAV-PHP.eB-GFP treated *ChAT-Cre;Tardbp^f/f^*mice became moribund. While all AAV-PHP.eB-GFP treated *ChAT-Cre;Tardbp^f/f^*mice reached end-stage disease around 47 weeks of age, all AAV-PHP.eB-CTR treated *ChAT-Cre;Tardbp^f/f^* mice remained alive to at least 110 weeks of age (**Fig. 2h**). Importantly, none of these animals displayed evidence of limb paralysis, unlike AAV-PHP.eB-GFP treated *ChAT-Cre;Tardbp^f/f^*mice (**Extended Data Fig. 6**). These results establish that the BBB permeable AAV-PHP.eB-CTR restores TDP-43 function in motor neurons and rescues disease phenotypes when delivered intravenously after onset of symptoms in a mouse model that mimics the early phase of ALS. These findings strongly support CTR as a promising AAV gene therapy designed to target TDP-43 dysfunction in ALS.

### Rescue of motor neuron disease correlates with prevention of cryptic exon inclusion by AAV-PHP.eB-CTR

To determine whether the striking rescue of muscle strength, body weight loss and premature death of *ChAT-Cre;Tardbp^f/f^* mice by AAV-PHP.eB-CTR was correlated with cryptic exon repression and motor neuron survival, we performed a full transcriptomic analysis of the lumbar spinal cord to see the effect of AAV-PHP.eB-CTR on TDP-43 associated mis-splicing. We see that multiple cryptic exon targets (*Mical2*, *Cx3cl1* and *Synj2bp*) associated with TDP-43 loss in the spinal cord tissue from AAV-PHP.eB-GFP treated *ChAT-Cre;Tardbp^f/f^* mice are rescued in *ChAT-Cre;Tardbp^f/f^*mice treated with AAV-PHP.eB-CTR (**Fig. 3a-c**). To corroborate this finding in motor neurons, we then used an integrated co-detection assay to monitor both the inclusion of cryptic exons in combination with the presence of ChAT, mouse endogenous TDP-43 or CTR. Consistent with previous RNA-seq analyses^3–7, 18^, transcripts of these cryptic exons served as molecular markers of TDP-43 loss of function (**Fig. 3d**). We found that CTR expression prevented inclusion of these cryptic exons in both spinal (**Fig. 3e,g**) and facial (**Fig. 3f,g**) motor neurons of AAV-PHP.eB-CTR treated *ChAT-Cre;Tardbp^f/f^*mice. Motor neurons of AAV-PHP.eB-CTR treated *ChAT-Cre;Tardbp^f/f^*mice which express CTR do not display inclusion of cryptic exons while non-rescued neurons continue to express cryptic exons, indicating the specific rescue of TDP-43 splicing function (**Fig. 3e,f**). Triple immunofluorescence staining of spinal cord sections demonstrated CTR expression in ChAT-positive neurons with nuclear depletion of mouse endogenous TDP-43 (**Extended Data Fig. 3**), further corroborating the co-detection assay and supporting the cryptic exon rescue potential of CTR in surviving motor neurons (**Fig.3e**). The RT-PCR analysis using primers against selected cryptic exon targets (*Mical2* and *Synj2bp*) and Basescope-ISH RED assay using probes targeting selected cryptic exons (*Ift81*, *Synj2bp* and *Unc13a*) also confirmed the rescue in AAV-PHP.eB-CTR treated *ChAT-Cre;Tardbp^f/f^*mice compared to AAV-PHP.eB-GFP treated *ChAT-Cre;Tardbp^f/f^* mice (**Fig.3h-j, Ext Data Fig. 7**). These results establish that the intravenous delivery of BBB permeable AAV-PHP.eB-CTR restores TDP-43 splicing repression function in motor neurons after symptom onset in a mouse model that mimics the early phase of ALS.

**Fig. 3:**
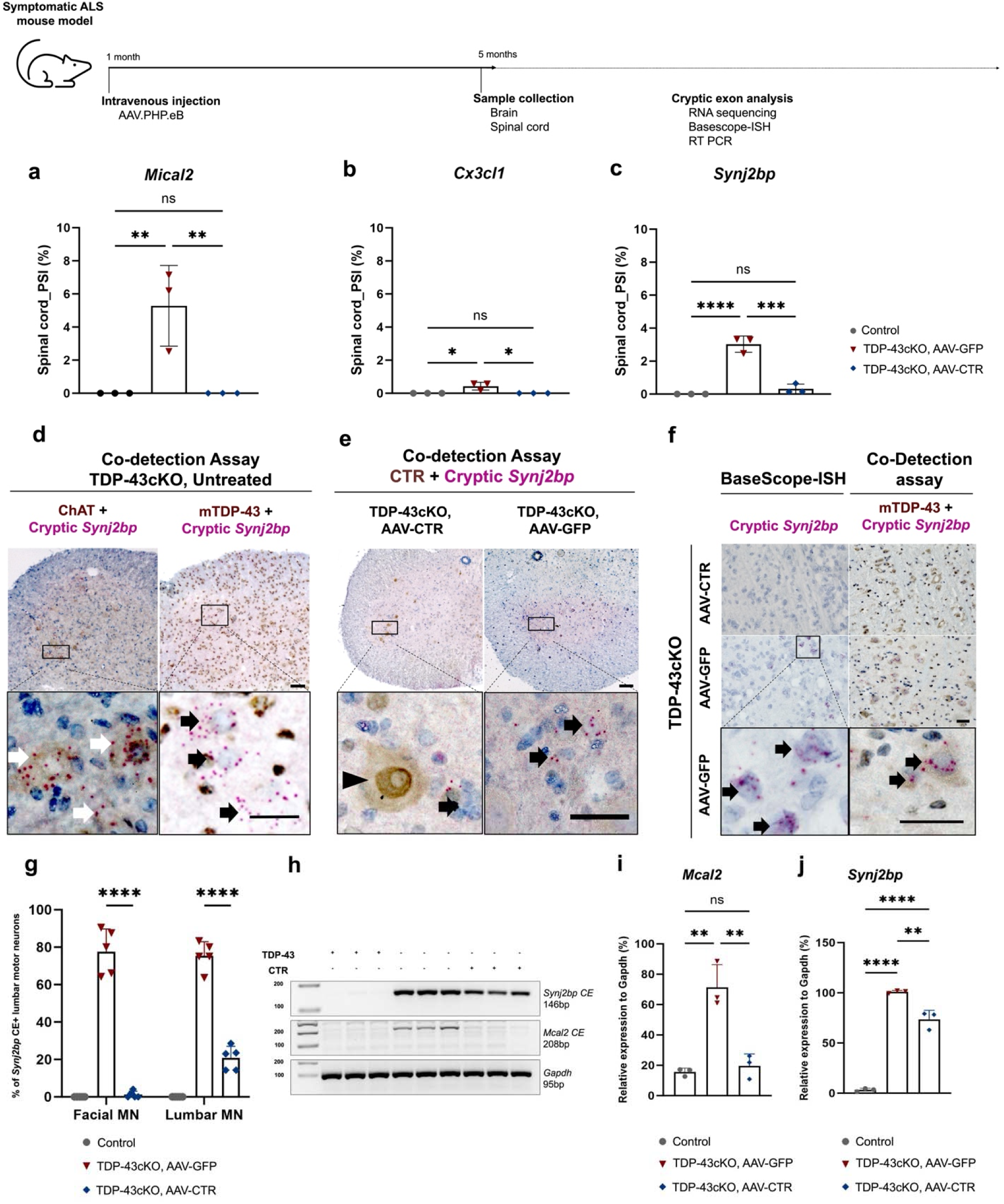
AAV-PHP.eB-CTR prevents the inclusion of TDP-43 regulated cryptic exons. (a-c) PSI values obtained from the RNA-seq analysis performed on lumbar spinal cord tissues from 5-month-old mice indicate the presence of cryptic exons in *Mical2*, *Cx3cl1,* and *Synj2bp*. This is rescued in the TDP-43cKO mice treated with AAV-PHP.eB-CTR (n=3 per group) **(d)** Representative images of simultaneous detection of the cryptic exon in *Synj2bp* using BaseScope-ISH probe and antibodies against ChAT or mouse endogenous TDP-43 reveals an elevated presence of transcripts containing *Synj2bp* cryptic exon in some ChAT positive neurons or in neurons lacking mouse endogenous TDP-43 in TDP-43cKO mice. Scale bar=100 µm. White arrows indicate *Synj2bp* cryptic exon in ChAT positive neurons. Black arrows indicate *Synj2bp* cryptic exon in TDP-43 deficient cells. Scale bar=20 µm. (**e**) Representative images of co-detection of cryptic exon in *Synj2bp* with CTR (using anti-human TDP-43) shows the rescue of cryptic exon expression in CTR expressing cells in TDP-43cKO mice. Scale bar=100 µm. Arrowhead indicates CTR expressing motor neuron, and black arrow shows CTR or mouse endogenous TDP-43 deficient cell expressing transcripts containing *Synj2bp* cryptic exon. Scale bar=20 µm. (**f**) Representative images from BaseScope-ISH or Co-detection assay to detect *Synj2bp* cryptic exon in facial motor nuclei. Scale bar=50 µm. Arrows depict TDP-43 deficient cells containing transcripts with *Synj2bp* cryptic exons in AAV-PHP.eB-GFP-treated TDP-43cKO mice. Scale bar=50 µm. (**g**) Quantitative analysis of ChAT positive motor neurons expressing *Synj2bp* cryptic exon in 5-month-old controls and TDP-43cKO mice treated with AAV-PHP.eB-GFP or AAV-PHP.eB-CTR (n=5 per group). (**h-j**) RT-PCR analysis of selected cryptic exons in targets such as *Mical2* and *Synj2bp* in 5-month-old controls and TDP-43cKO mice treated with AAV-PHP.eB-CTR or AAV-PHP.eB-GFP (n=3 per group).

### AAV.PHP.eB-CTR attenuates motor neuron loss, prevents ventral root degeneration and rescues denervation muscle atrophy in *ChAT-Cre;Tardbp^f/f^*mice

To determine whether prevention of TDP-43 cryptic exons led to attenuation of motor neuron loss, we quantified the number of ChAT positive motor neurons in 5-month-old *ChAT-Cre;Tardbp^f/f^* mice treated with AAV-PHP.eB-CTR and AAV-PHP.eB-GFP. We observed loss of ∼50% of motor neurons in AAV-PHP.eB-GFP-treated *ChAT-Cre;Tardbp^f/f^* mice compared to littermate controls (**Fig. 4a-d**). In contrast, AAV-PHP.eB-CTR significantly attenuated the loss of these cells (**Fig. 4a-d**). Concordant with the rescue of motor neurons in the *ChAT-Cre;Tardbp^f/f^*mice treated with AAV-PHP.eB-CTR, both light microscopy and TEM analyses showed restoration of large diameter fibers in the ventral roots (**Fig. 4e-h**) and marked attenuation of skeletal muscle atrophy in the hindlimbs (**Fig. 4i & Extended Data Fig. 8**). These results strongly indicate that the rescue of motor neuron disease was conferred by the robust protection of motor neurons due to CTR-mediated repression of TDP-43-regulated cryptic exons within ∼80% of motor neurons. Our findings validate this AAV-based strategy using CTR to restore multiple cryptic targets of TDP-43, offering a novel, mechanism-based gene therapy for the treatment of ALS.

**Fig. 4:**
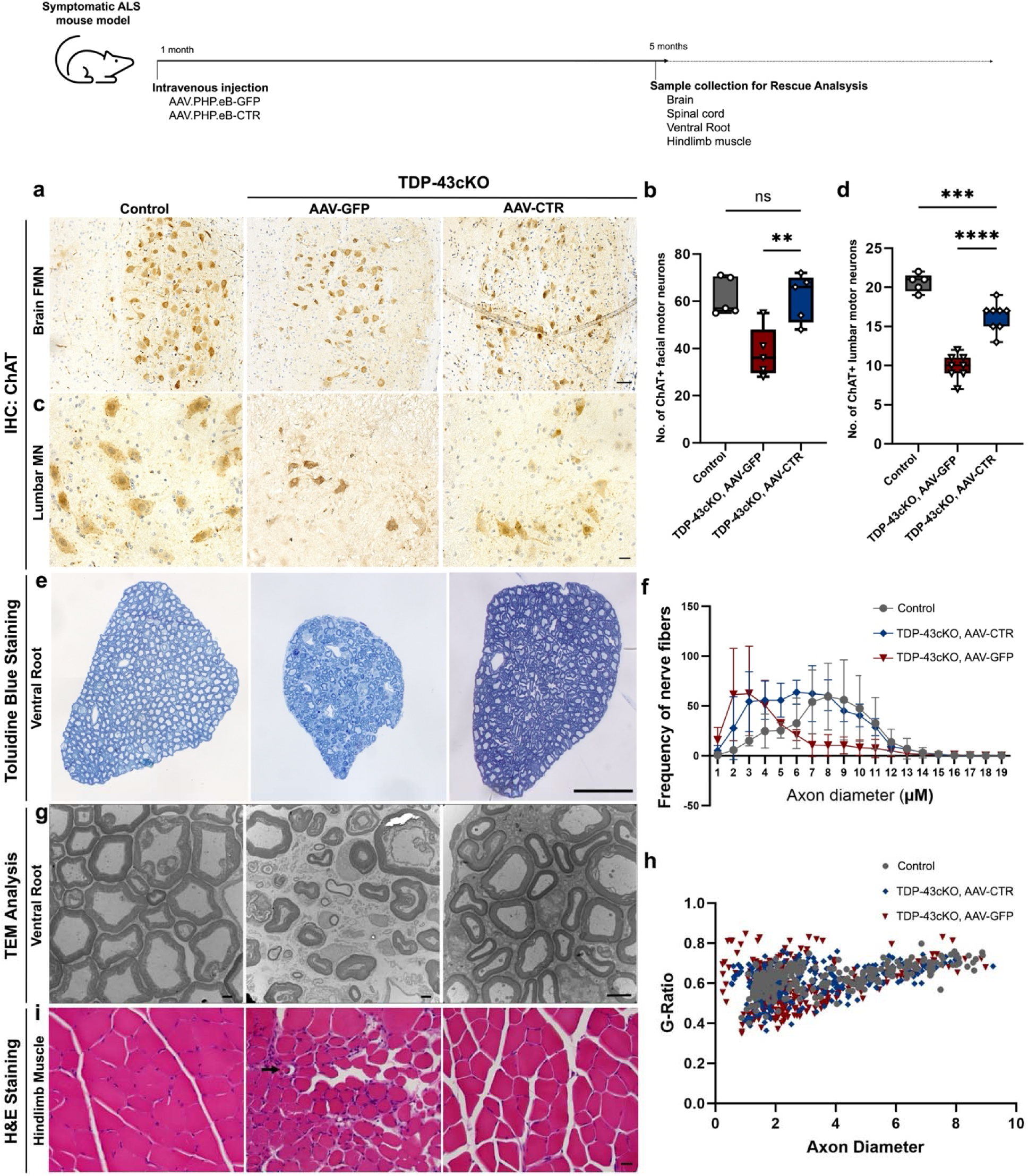
AAV-PHP.eB-CTR attenuates motor neuron loss, axon degeneration, and skeletal muscle atrophy. (**a, c**) Representative images from pathological analysis in facial (**a,** Scale bar=50 µm) and spinal (**c,** Scale bar=20 µm) motor neurons showing that CTR protects against motor neuron loss in adult TDP-43cKO mice treated with AAV-PHP.eB-CTR at symptomatic stage (treated at 1.5 months of age and analyzed at 5 months of age, n=8). (**b, d**) Quantification of number of ChAT positive motor neurons in facial motor nuclei (**b**, p<0.05, n=5) and the lumbar spinal cord (**d**, p<0.05, n=8). (**e**) Representative toluidine blue-stained light microscopic images of transversely sectioned motor roots (Scale bar=50 µm). (**f**) A notable reduction in the number of large axonal fibers is observed in the TDP-43cKO mice treated with AAV-PHP.eB-GFP compared to those treated with AAV-PHP.eB-CTR (p<0.05, n=3 per group). (**g, h**) Representative TEM images and G-ratio plot of transversely sectioned roots which shows the rescue of the degeneration of axonal fibers in TDP-43cKO mice treated with AAV-PHP.eB-CTR compared to the TDP-43cKO mice treated with AAV-PHP.eB-GFP (n=3 per group). Scale bar=2 µm. (**i**) Representative images after H&E staining of gastrocnemius muscle sections (10 µm) from control mice and TDP-43cKO mice treated with AAV-PHP.eB-CTR or AAV-PHP.eB-GFP at 5 months of age. Arrow represents the muscle fiber undergoing degeneration in TDP-43cKO mice treated with AAV-PHP.eB-GFP, which is rescued in TDP-43cKO mice treated with AAV-PHP.eB-CTR (Scale bar=50 µm).

### As compared to brain neurons, human and mouse spinal motor neurons express lower level of *UNC13A*

Since preventing cryptic splicing of *Unc13a* is sufficient to attenuate cognitive decline in mice lacking TDP-43 in brain neurons (personal communication), but was insufficient to attenuate motor neuron disease, we asked whether the failure could be explained by the lower levels of *Unc13a* occurring in spinal motor neurons. We compared Unc13a protein levels between the brain and spinal cord of wildtype mice and found a markedly lower level in spinal cord than in the brain (**Fig. 5a, b).** Moreover, the comparison of the level of *Unc13a* cryptic exon expression in both cortical and spinal neuron-specific TDP-43 knockouts revealed a higher expression in cortical neuron knockouts **(Ext Data Fig. 9a**). RNA-sequencing in our cohort also failed to identify the presence of *Unc13a* cryptic exon in the spinal cord samples from the TDP-43cKO mice (**Ext Data Fig. 9b**).

**Fig. 5:**
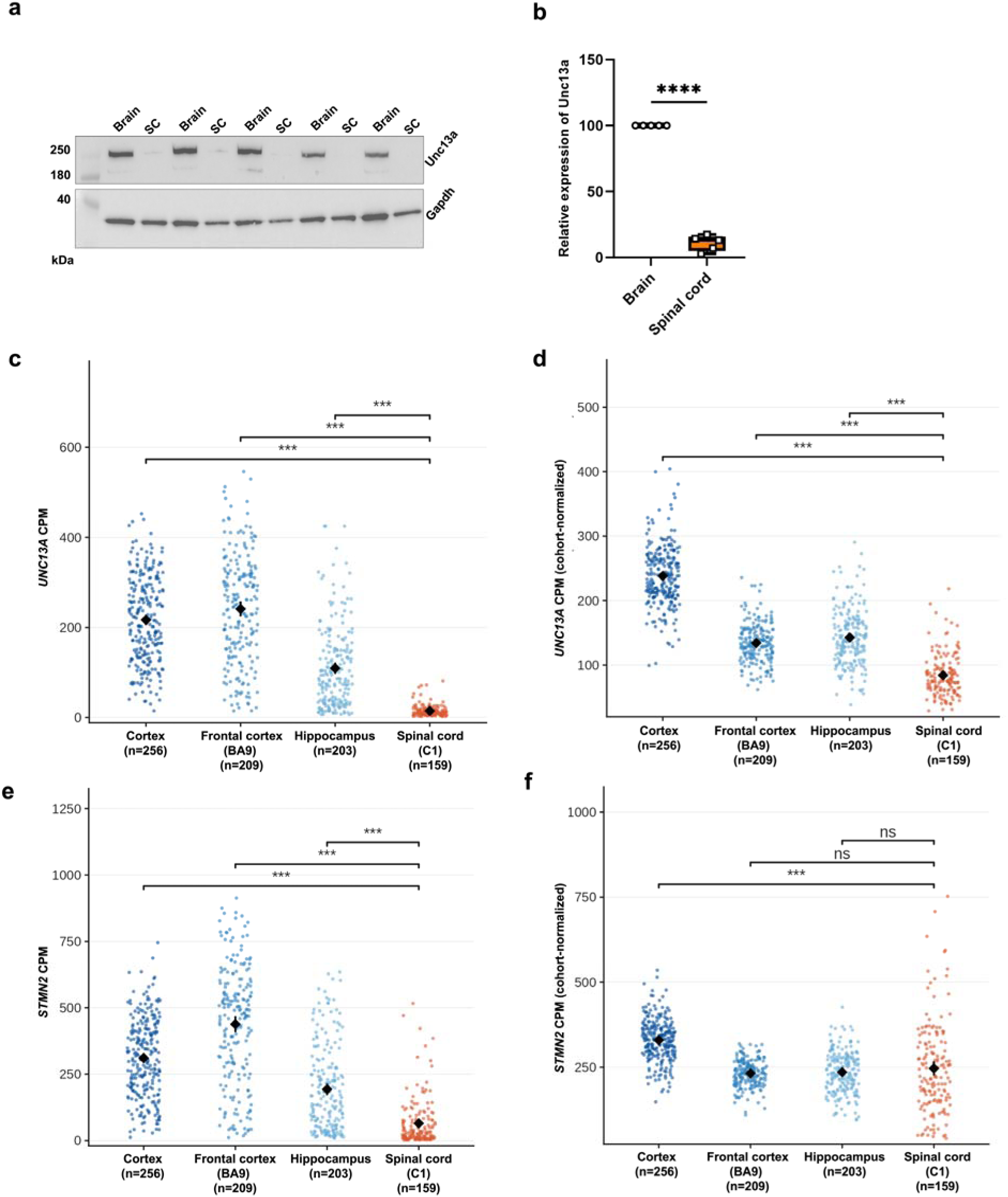
(**a, b**) Immunoblot analysis of protein extracts from 1-month-old wildtype mice showed that the level of Unc13a protein is significantly lower in the spinal cord as compared to the brain (p<0.0001; n=5). (**c**) Raw *UNC13A* expression (counts per million, CPM) in four human CNS regions from the Genotype-Tissue Expression (GTEx) project (Cortex, n=256; Frontal cortex BA9, n=209; Hippocampus, n=203; Spinal cord (cervical C1), n=159. Gene-level counts accessed via the Snaptron/recount3 GTExcompilation). Each point is one donor sample; black diamond and error bars show mean ± 95% CI. *UNC13A* CPM was significantly lower in spinal cord than in cortex, frontal cortex BA9, and hippocampus (two-sided Wilcoxon rank-sum test vs. spinal cord; p<0.0001 for each; *** p<0.001, ** p<0.01, * p<0.05, ns = not significant). (**d**) *UNC13A* CPM normalized to each sample’s estimated relative neuronal content, in the same GTEx samples as (**c**). (**e**) Raw *STMN2* expression (CPM) in the same GTEx samples as in panel c. *STMN2* CPM was significantly lower in spinal cord than in cortex, frontal cortex BA9, and hippocampus (p<0.0001 for each). (**f**) *STMN2* CPM normalized to estimate relative neuronal content, as in panel d.

To determine the level of *UNC13A* expression in human brain and spinal cord, we analyzed publicly available bulk RNA-seq data. Analysis of human GTEx bulk RNA-seq data showed that *UNC13A* transcript levels were markedly lower in spinal cord as compared to each of brain regions examined, both as raw counts per million (CPM) (**Fig. 5c**) and after correcting each sample’s CPM for its neuronal content using the marker-gene deconvolution method BRETIGEA^34^. Normalized *UNC13A* was significantly higher in cortex (3.0-fold), frontal cortex (1.7-fold), and hippocampus (1.8-fold) than the spinal cord (p<0.0001) (**Fig. 5d**). The persistence of this difference after accounting for the lower overall neuronal content of bulk spinal cord tissue indicates that *UNC13A* is expressed at a lower level per neuron in spinal cord than in other regions, consistent with single-nucleus data showing that *UNC13A* is specifically and markedly reduced in spinal alpha-motor neurons relative to other neuronal populations^3^. While raw *STMN2* CPM was also substantially lower in spinal cord as compared to the brain (**Fig. 5e**), normalized *STMN2* remained significantly higher in the cortex (1.5-fold, p<0.0001) but was no longer significantly different between spinal cord and frontal cortex (p=0.80) or hippocampus (p=0.68) (**Fig. 5f**), which implies that the reduction in *STMN2* in spinal cord largely reflects the lower neuronal content. This region-specific enrichment of *UNC13A* offers a likely explanation for the insufficiency of restoring *Unc13a* alone to attenuate motor neuron disease in mice. Furthermore, it is important to note that, as *UNC13A* is also expressed at lower levels in human spinal cord tissue, it is also possible that the selective vulnerability of brain neurons to *UNC13A* levels may occur in human disease, including FTLD-TDP, LATE and AD.

## DISCUSSION

In view of the nonconserved nature of cryptic exons associated with TDP-43 dysfunction^7^, it is expected that those found in humans should not overlap with those in mice. As such, sequences targeted by TDP-43 for splicing regulation in humans should differ from those in mice. However, it is possible that certain gene targets can still be shared between divergent species, and *UNC13A* is one such example. In this case, the cryptic exon in humans is localized to intron 20^3,23^-which is distinct from the cryptic exon found in intron 1 of mouse *Unc13a*^18^. Coincidentally, both cryptic exons introduce a premature termination codon, leading to nonsense mediated decay of the pre-mRNA^3,4,7^. This unexpected observation provides an opportunity to determine whether the exclusion of *Unc13a* cryptic exon alone is sufficient to mitigate motor neuron disease.

Genetic variants in *UNC13A* which facilitate higher levels of its cryptic transcripts in the context of TDP-43 depletion in ALS-FTD and AD-TDP cases^3,4^ have strongly encouraged antisense oligonucleotide (ASO) therapeutic strategies designed to correct *UNC13A* cryptic splicing in ALS^15,16,24^. However, direct evidence supporting the sufficiency of solely preventing *UNC13A* cryptic splicing to mitigate motor neuron disease is lacking. In this study, we leveraged the shared consequence of TDP-43-associated cryptic exon inclusion in the mouse and human homologues of the gene, *UNC13A* to investigate the sufficiency of preventing the *Unc13a* cryptic exon in rescuing motor neuron disease.

We first validated that genetic ablation of the cryptic exon of *Unc13a* in mice lacking TDP-43 normalized the expression levels of *Unc13a* (personal communication). To determine whether the exclusion of *Unc13a* cryptic exon was independently sufficient to influence motor neuron disease, we analyzed *ChAT-Cre;Tardbp^f/f^* mice lacking the *Unc13a* cryptic exon but found no evidence of attenuation of motor neuron disease (Fig. 1), suggesting that *Unc13a* may not play a major role in spinal motor neuron loss and restoration of additional cryptic targets may be necessary for rescue. Supporting this notion is the rescue of motor neuron disease in the AAV-PHP.eB-CTR treated *ChAT-Cre;Tardbp^f/f^* mice in which multiple TDP-43 cryptic targets, including *Unc13a*, are normalized (Figs. 2–4).

Since the exclusion of the *Unc13a* cryptic exon is sufficient to preserve cognition in a mouse model lacking TDP-43 in forebrain neurons (personal communication), we surmise that the contribution of cryptic *Unc13a* plays a major role in brain neurons than spinal cord. We speculate that one factor to account for such vulnerability of brain neurons to cryptic *Unc13a* is the regional specificity of gene expression. Indeed, we found that the level of *Unc13a* in the spinal cord as compared to that of the brain (Fig. 5) is significantly lower, consistent with the view that *Unc13a* plays a greater role in the brain. We thus propose that targeting *UNC13A* cryptic exon may not be sufficient for spinal motor neurons but holds promise for TDP-43 proteinopathy associated with cortical regions, such as FTD-TDP. We further speculate that cryptic *STMN*2 may play a dominant pathogenic role in spinal motor neurons^6,24^, and strategies designed to correct cryptic splicing of *STMN2* hold promise to mitigate spinal motor neuron loss in ALS. To achieve optimal clinical outcomes, we suggest that it may be necessary to correct multiple cryptic targets to mitigate loss of upper and lower motor neurons. As such, strategies designed to target both *STMN2* and *UNC13A* for TDP-43 dysfunction hold greater promise for ALS^25^.

In contrast to neonatal delivery of CTR using AAV9^7,19^, this study provides proof of concept for systemic delivery of CTR using a BBB-penetrant AAV at early symptomatic stage to prevent TDP-43 cryptic splicing, preserve motor neuron integrity, prevent limb paralysis and maintain normal lifespan supports the view that restoration of multiple cryptic targets is predicted to provide optimal clinical benefit, offering broader therapeutic potential for ALS. The emerging need for such a strategy to restore multiple cryptic targets is emphasized by discoveries of additional cryptic targets of TDP-43 as revealed by studies in which NMD is inhibited, indicating widespread transcriptome and proteome effects^16,17^. While AAV-PHP.eB is a robust BBB-crossing vector in mouse models and transduced >70% of spinal motor neurons to deliver CTR and prevent motor neuron disease, this serotype is not permeable to the human BBB. Emerging next generation AAV serotypes with the potential to cross the human BBB^26^ will provide a strong foundation for the clinical translation of our mechanism-based gene therapy designed to restore multiple TDP-43 cryptic targets, including *STMN2* and *UNC13A*. Such advances hold promises not only for ALS, but also for other neurodegenerative disorders associated with TDP-43 proteinopathy, including FTLD-TDP^8^, LATE^27^, and AD-TDP^28–31^.

## MATERIALS AND METHODS

### Animals

Our conditional *Tardbp* knockout mice (*Tardbp^f/f^*, Jax Stock 017591) in which exon 3 is flanked by LoxP sites^32^ were crossbred with ChAT-dependent Cre driver line (*ChAT-Cre)* transgenic mice on a C57BL/6J background (Jax Stock 006410) to generate a cohort of *ChAT-Cre;Tardbp^f/+^*mice. These heterozygous mice were then interbred to generate *ChAT-Cre/ChAT-Cre;Tardbp^f/+^*, which was subsequently bred with *Tardbp^f/f^* mice to generate a final cohort of *ChAT-Cre;Tardbp^f/+^* (control mice) and *ChAT-Cre;Tardbp^f/f^*(TDP-43 conditional knockout mice) (**Extended Data Fig.1**). Animals were housed in a 12-hour light/dark cycle with food and water *ad libitum*. All experiments were performed in accordance with the National Institute of Health (NIH) Guidelines and approved by the Animal Care and Use Committee (ACUC) at Johns Hopkins Medicine.

The *Unc13a* cryptic exon (CE) deletion mouse model was generated using CRISPR-Cas9 technology. Specifically, a ∼44 bp genomic region corresponding to the CE was deleted, producing *Unc13a CE^-/-^* mice at an in-house transgenic core facility at Johns Hopkins Medicine. This deletion was confirmed by PCR amplification using primers flanking the CE region (Forward: GGCTCTCCTGTCTTCTCAGC and Reverse: CTTTGTGTGCAAGGCACGAA). To evaluate the effect of the *Unc13a* cryptic exon in the context of TDP-43 depletion, *Unc13a CE^-/-^* mice were crossed with *Tardbp^f/f^*;*ChAT-Cre* mice to generate the final cohort of *ChAT-Cre;Tardbp^f/f^*:*Unc13aCE^-/-^*mice (hereafter referred to as TDP-43cKO;*Unc13aCE^-/-^*) which is deficient in both the *Unc13a* cryptic exon and TDP-43 within the motor neurons.

### Experimental Design

The male or female littermates including control and TDP-43cKO mice from the final breeding cages were randomly divided into three groups; 1) *ChAT-Cre;Tardbp^f/+^*mice treated with either AAV-PHP.eB-CTR or AAV-PHP.eB-GFP (n=20); 2) *ChAT-Cre;Tardbp^f/f^*mice treated with AAV-PHP.eB-GFP (n=20); and 3) *ChAT-Cre;Tardbp^f/f^*mice treated with AAV-PHP.eB-CTR (n=20). For additional analysis, litters from the same breeding pairs were included to analyze the rate of AAV-PHP.eB transduction efficiency at 1-, 3-, and 5-month of age, and to perform cryptic exon and RNA analysis.

### Viral Vector Packaging and Delivery

AAV-PHP.eB packaging was performed by Virovek (Hayward, CA). The expression of CTR or GFP was confirmed independently by HeLa cell transduction and protein blot analysis prior to all experiments (data not shown). Intravenous delivery of AAV-PHP.eB vector was performed through lateral tail vein injection in 6-week-old mice. Briefly, mice were warmed using an overhead heat lamp for 2-3 minutes to dilate the veins. The animal was lightly anesthetized and restrained using a commercial device (Tail veiner Restrainer, Braintree Scientific, Inc.). The AAV-PHP.eB vector solution was loaded into a 1 mL syringe using a 30-G needle without air bubbles. The tail was grasped at the distal end using the index and middle fingers while keeping the lower part of the tail held between the thumb and the ring fingers. The needle entered the vein at a shallow depth while maintaining the syringe and needle parallel to the tail. A total of 50 mL vector solution (1E+12 vg/mL, from stock 2E+13 vg/mL) diluted in 100 mL sterile PBS was injected when a flash of blood was detected in the hub of the needle at the time of insertion without any resistance. After administration, the needle was removed, and gentle compression was applied until bleeding stopped. The animals were returned to their cages after recovering from the anesthesia.

### Grip Strength Test

Grip strength analysis for both forelimbs and hind limbs was performed using a metal grid in a blinded manner. Briefly, the animals were placed on the center of a wire mesh cage top and inverted in such a way that only the forelimb and hindlimb paws could grasp the mesh. To perform the test, the grid was shaken gently enough for the animal to hold it and then turned upside down over an empty cage. The latency to fall (the duration each mouse was able to hang upside down) was measured in 3 trials for each animal with a maximum duration of 1 min. The mice were given ∼2 min interval between the attempts and the best attempt was used for the grip strength analysis.

### Survival Analysis

The body weight and overall health of all animals were monitored daily during the whole experiment to monitor disease progression. Upon hindlimb paralysis, mice were provided wet chow and Dietgel. End stage was defined by the loss of the righting reflex, i.e., failure of the mouse to right itself within 10 seconds when placed on its back. At this stage, the mice were euthanized, and tissue samples were collected for further analysis.

### Tissue Collection

Mice were euthanized and transcardial perfusion was done with ice-cold phosphate buffered saline (1X PBS, 100 mL) and then with 4% paraformaldehyde (PFA, 50mL, freshly prepared in 1X PBS). The brain and spinal cord (divided into cervical, thoracic, and lumbar regions) were dissected out, post-fixed overnight in ice-cold 4% PFA and then processed for paraffin embedding and sectioning. To extract the RNA, whole spinal cords were subjected to hydraulic extrusion using a 1 mL syringe and a 20-gauge needle after perfusion with 1X PBS. The tissues were flash frozen and stored at –80°C until further analysis.

### Histology and Immunohistochemistry

Formalin-fixed paraffin embedded sections were used for histology and immunohistochemistry analysis. 10 µm thick coronal sections of spinal cord were cut and stained with Cresyl violet (Nissl stain to label the total population of cells and show the cellular distribution in the area under investigation) for histological analysis. For immunohistochemistry, sections were de-paraffinized, subjected to antigen retrieval in 10mM sodium citrate buffer at 100°C for 5min, and blocking (1.5% normal goat serum and 0.1% Triton-X in PBS) at room temperature for 1 hour. The sections were then incubated overnight at 4°C with primary antibodies diluted in blocking buffer. The following primary antibodies were used, ChAT (Cat#AMAb91130, 1:1000, Atlas Antibodies), human-specific TDP-43 (Cat#WH0023435M1, 1:1000, Millipore Sigma) and mouse-specific TDP-43 (C-Terminus, Cat#12892-1-AP, 1:1000, Proteintech). CTR was detected using the human-specific TDP-43 antibody. Then, the sections were washed three times with 1X PBS containing 1% Tween-20 (PBST) followed by incubation in biotinylated secondary antibodies (Anti-Mouse IgG, Cat#BP-9200 and Anti-Rabbit IgG, Cat#BP-9100, Vector laboratories) diluted in blocking buffer at room temperature for 1 hour. Sections were washed three times with PBS-T and incubated with avidin-biotin peroxidase complex kit (Cat#PK-7100, Vectastatin ABC Reagent, Vector Laboratories) for 1 hour. After the incubation, the sections were washed three times with PBST and incubated with DAB solution for 30 seconds to 1 minute to produce brown reaction product. The sections were counterstained with hematoxylin, dehydrated and mounted. The images were acquired using Zeiss Apotome Inverted Brightfield Microscope (Zeiss, Germany) and quantification of motor neurons was done using ImageJ software (National Institutes of Health, Bethesda, MD) from four serial sections per animal in a genotype and treatment-blinded manner independently by three investigators. The average number of total motor neurons was determined as normalized values to the total number of sections used.

### Immunofluorescence and Confocal imaging

The lumbar spinal cord sections were deparaffinized and rehydrated as described previously and blocked with 5% BSA blocking buffer. For triple immunofluorescence staining, sections were incubated with following primary antibodies: goat anti-ChAT antibody (Cat#NBP1-30052, Novus Biologicals), mouse anti-TARDBP monoclonal antibody [clone: 2E2-D3] (Cat#H00023435-M01, Novus Biologicals) and rabbit anti-TDP-43 polyclonal antibody (C-Terminal) (Cat#12892-1-AP, ProteinTech) that were incubated overnight at 4°C. Following primary antibodies, the sections were washed and incubated for 1hour at RT with respective fluorophore-conjugated secondary antibodies (1:400) including donkey anti-mouse 594 (Cat#A-11001, Thermofisher), Donkey anti-goat 647 (Cat#A-21450, Invitrogen) and Donkey anti-rabbit 488 (Cat#A32790, Invitrogen). Images were acquired using Mica Confocal microscope at 63x magnification (Leica Microsystems).

### *In Situ* Hybridization: BaseScope^TM^ RED and Co-detection Assay

RNA *in situ* hybridization was performed using BaseScope Detection Reagent v2-RED Assay Kit (Cat#323900, Advanced Cell Diagnostics, ACDBio, USA) following the manufacturer’s instructions. 3ZZ probes were designed targeting the cryptic exon sequences within mouse *Synj2bp* (BA-Mm-Synj2bp-E2-intron2-NJ, Cat#712191, ACDBio, USA), *Ift81* (BA-Mm-Ift81-E17-intron17-NJ, Cat#712201, ACDBio, USA) and *Unc13a* (BA-Mm-Unc13a-O1-2EJ-C1, Cat#1182491-C1, ACDBio, USA). To detect the infectivity of AAV, we have designed antisense probes against the AAV-PHP.eB vector backbone and performed BaseScope RED assay on spinal cord and brain tissue sections (BA-CMV-Enhancer-2zz-st-sense-C1, Cat#1039721-C1, ACDBio, USA). To determine the accumulation of CTR transcripts, we have also designed a 1zz BaseScope probe against the CTR mRNA (BA-Hs-TARDBP-RAVER1-1zz-C1, Cat#1573691-C1, ACDBio, USA). Briefly, formalin-fixed paraffin-embedded sections (10 µm) were deparaffinized and pre-treated with hydrogen peroxide for 10 min at room temperature, target retrieval buffer for 15 min at 99°C and protease IV for 30 min at 40°C. The following day, sections were hybridized with the target probes in a HybEZ^TM^ II Hybridization Oven (ACDBio, USA) for 2 hours at 40°C. The hybridized signals were amplified through a series of amplification steps at 40°C and washing steps at room temperature. Finally, the signals were detected using a chromogenic red substrate and counterstained with hematoxylin. To assess the quality of RNA, each sample was hybridized with a probe targeting a housekeeping gene (BA-Mm-Ppib, Cat#701071, ACDBio, USA). As a control for the background staining, each section was also evaluated with another probe targeting a bacterial gene (BA-DapB, Cat#701011, ACDBio, USA). The co-detection assay was performed using the RNA-Protein Co-Detection Ancillary Kit (Cat#323180, ACDBio, USA) as per manufacturer’s instructions. The images were captured using a Zeiss Apotome Inverted Brightfield Microscope (Zeiss, Germany). The hybridized signal for each RNA transcript was detected as red punctate dots or clusters and quantified using ImageJ software (National Institutes of Health, Bethesda, MD). To compare the *Unc13a* cryptic exon level in the brain, we used tissue sections from *CaMKII-CreER;Tardbp^f/f^*mice in which TDP-43 is selectively depleted in forebrain neurons^33^.

### RNA Sequencing

The lumbar spinal cord tissues were homogenized using a Kimble Pellet Pestle Cordless Motor (Cat#749540, DWK Life Sciences) and total RNA was extracted using TRIzol (Life Tech., 15596-026) and RNeasy Mini Kits (Cat#74106, Qiagen). Total RNA for RNA-Seq was then processed using the TruSeq Stranded Total RNA Library Prep Kit (Illumina) to construct RNA-Seq libraries. Sample libraries were then sequenced on an Illumina NextSeq. Data was de-multiplexed and converted into fastq files. Fastq files were then processed by STAR^34^ and splice junction counts were used to calculate cryptic exon PSI^18^.

### RNA extraction and RT-PCR

The spinal cord tissues were homogenized using a Kimble Pellet Pestle Cordless Motor (Cat#749540, DWK Life Sciences) and total mRNA was extracted using RNeasy Mini kit (Cat#74106, Qiagen) following the manufacturer’s instructions. The extracted RNA was quantified using Nanodrop One (Cat#269-309101, Thermo Fisher Scientific, Inc.) and then converted to cDNA using the Protoscript II First Strand cDNA Synthesis Kit (Cat#E6550L, New England Biolabs Inc.). To detect the cryptic exon containing mRNA upon TDP-43 depletion, we PCR amplified cDNA using primers specifically designed to target the cryptic exon and another primer spanning the canonical exon. We used primers against *Mical2* (Forward-5’GCG AGA CCT TGG GTC AAG GAG-3’, Reverse-5’-GCA GGC ATT CTG AAG CTG TG-3’) and *Synj2bp* (Forward-5’-GGGTGATAAGATCCTCTCGGG-3’, Reverse-5’-TCTTCCTGAGGACCTCCGTT). *Gapdh* measurements were used for normalization. The resulting PCR products were then resolved on a 1.5% agarose gel containing GelRed Nucleic Acid Stain (Cat#SCT121, Millipore Sigma) and visualized under UV light to detect the presence of respective products, indicative of cryptic exon-containing mRNA. Images were analyzed using ImageJ software.

### Ventral Root Extraction and Light Microscopy Analysis

To extract the lumbar dorsal root ganglion (DRG) with their corresponding roots, animals were perfused with freshly prepared 2% glutaraldehyde, 2% paraformaldehyde (EM grade prill), 50 mM sodium cacodylate, 50 mM phosphate (Sorenson’s) 3 mM MgCl_2_, pH 7.2 at 1085 mOsm. After perfusion, animals were kept in the cold room (4°C) for two hours and then the DRGs with corresponding roots were extracted and incubated in fresh fixative overnight at 4°C. Samples were processed as follows: all incubations were carried out at 4°C until the 70% ethanol step, then incubated at room temperature. Initially, samples were rinsed in a buffer consisting of 75 mM cacodylate, 75 mM phosphate, 3.5% sucrose, 3 mM MgCl_2_, pH 7.2 with an osmolarity of 430 mOsm, for 45 minutes. Following the buffer rinse, samples were microwave-treated using a Pelco laboratory grade microwave model 3400, with a power setting of 50% and a pulse duration of 10 seconds, followed by a 20-second pause and then another 10-second pulse. The samples were then post-fixed in a solution containing 2% osmium tetroxide, 1.6% potassium ferrocyanide, and the same buffer without sucrose, and were placed on ice in the dark for 2 hours. Samples were then rinsed in 100 mM maleate buffer with 3.5% sucrose pH 6.2. This was followed by En-bloc staining of the samples for 1 hour with filtered 2% uranyl acetate in maleate sucrose buffer, pH 6.2. Samples were then dehydrated through a graded series of ethanol to 100%, transferred through propylene oxide, and embedded in Eponate 12 (Pella). The samples were then cured at 60°C for 2 days.

Each spinal ganglion with roots was embedded in epoxy resin, and sagittal sections were cut on Reichert Ultracut E Microtome with a Diatome Diamond knife (450) to get 1 μm semithin sections. Toluidine blue staining was performed on each section to visualize the myelinated nerve fibers and images were acquired using a Zeiss Apotome Inverted Brightfield Microscope (Zeiss, Germany). Three animals per group were analyzed to quantify nerve fibers. The quantifications were performed independently by three investigators in a blinded manner to the genotype and treatment.

### Ventral Root: TEM analysis

To perform Electron Microscopy, 60nM sections were cut and picked up on formvar coated 1X2mm copper slot grids and stained with methanolic uranyl acetate followed by lead citrate. Grids were viewed on a Hitachi 7600 Transmission Electron Microscope (TEM) operating at 80 kV and Digital images were captured with an XR50 5-megapixel CCD by AMT. Three animals per group were analyzed to quantify the nerve fibers and each section was analyzed using magnifications up to X5000. Around 100 fibers per animal were quantified independently by the investigators in a blind manner to the genotype and treatment.

### Muscle Histology

The hind limb muscles were dissected, and their gross morphology and weight were recorded. Each muscle was fixed in 4% PFA overnight, and 10 mm paraffin embedded coronal sections were cut. The sections were deparaffinized and stained with hematoxylin and eosin (H&E). Myofiber morphology and diameter were assessed using a Zeiss Apotome Brightfield microscope (Zeiss, Germany).

### Immunoblotting

Brain tissue was dissected out, and whole spinal cord was flushed out using a 1mL syringe and 20-gauge needle and flash frozen until further analysis. Tissues were then homogenized (The Scilogex D160 Homogenizer, Marshall Scientific) in ice-cold lysis buffer containing protease inhibitor cocktail (Cat#78430, Thermo scientific), centrifuged at 4°C, 12,000 rpm for 20 min. The supernatant was collected and the protein concentration was determined through BCA Assay using Pre-diluted Protein assay standards (Cat#23208, Thermo scientific). 20 mg protein was resolved in NuPAGE 4-12% Bis-Tris gel (Cat#NP0323BOX, Invitrogen) along with Precision Plus Protein Dual Color Standards (Cat#1610374, Biorad) and transferred to a polyvinylidene difluoride membrane. The membrane was blocked with 5% non-fat milk in Tris-buffered saline with 1% Tween-20 (TBST) for 1 hour and then probed with primary antibody against Unc13a (1:2000, Cat# 55053-1-AP, Proteintech) and Gapdh (1:10000, Cat# 60004-1-IgProteintech). The detection was performed with respective horseradish peroxidase-linked secondary antibodies (Goat anti-rabbit IgG, HRP-linked Antibody Cat#7074, Cell Signaling Technology and Goat anti-Mouse IgG (H+L) Secondary Antibody, HRP, Cat#31430, ThermoFisher Scientific) and enhanced chemiluminescence detection system as per manufacturer’s instructions (Immobilon Western Chemiluminescent HRP Substrate, Cat#WBKLS0500, Millipore Sigma). As an internal control to normalize the protein levels, the blots were stripped and incubated with Gapdh primary antibody (1:10000). The images were taken using the ChemiDoc imaging system (Bio-Rad) and analyzed using ImageJ.

### GTEx data analysis

*UNC13A* and *STMN2* expression were analyzed in bulk RNA-seq data from the Genotype-Tissue Expression (GTEx) project, accessed as gene-level read counts via the Snaptron/recount3 GTEx compilation. Samples were restricted to CNS tissues passing quality control (RNA integrity number ≥5.5, “RNASEQ” analysis freeze, ≥5×10 uniquely mapped reads where available), yielding 2,661 samples across 13 CNS regions; four regions are shown here (Cortex, n=256; Frontal cortex BA9, n=209; Hippocampus, n=203; Spinal cord [cervical C1], n=159, the reference tissue). Raw expression was expressed as counts per million (CPM) using each sample’s total gene-level read coverage as the library-size denominator. Relative Neuronal content was estimated per sample using BRETIGEA^35^, a marker-gene-based deconvolution method for human brain bulk tissue: a neuronal surrogate proportion variable (SPV) was computed from the top 50 neuronal marker genes (SVD method), shifted to strictly positive values, and rescaled so the cohort mean equals 1; CPM was divided by this relative-content value. Two-sided Wilcoxon rank-sum tests compared each brain region to spinal cord, without correction for multiple comparisons. Code and data are available at https://github.com/caotianyu0427/gtex-unc13a-stmn2-bretigea.

### Statistical Analysis

Data analysis was performed through the following statistical tests using the Graphpad Prism v.5. software (GraphPad Software Inc., San Diego, CA, USA). Histological data was analyzed using an unpaired, two-tailed Student’s t-test with Tukey’s multiple comparison test and one-way analysis of variance (ANOVA) to identify significant differences between groups. Behavioral data was analyzed using two-way ANOVA with Tukey’s multiple comparison test. The Kaplan-Meier survival data were analyzed using log-rank test. The p-values less than 0.05 considered significant.

## EXTENDED DATA

**Extended Data Fig. 1:**
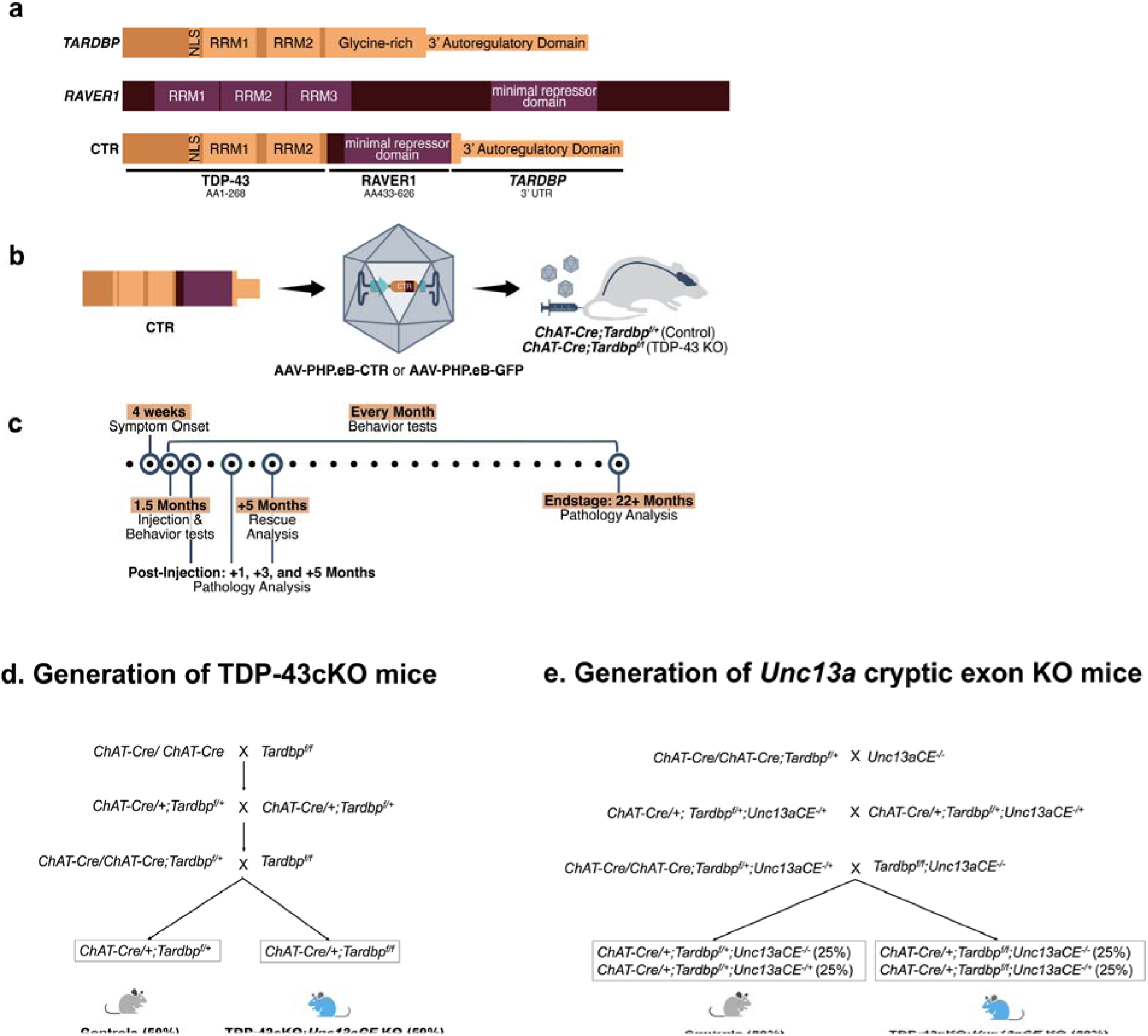
Schematic representations of structure of TDP-43, RAVER1, and CTR, CTR therapeutic strategy and study design. (**a**) The TDP-43 protein contains an N-terminal domain harboring a nuclear localization signal (NLS), two RNA-recognition motifs (RRMs), and a C-terminal domain which is glycine-rich and aggregation-prone. Parts of the final exon and 3’UTR encode an autoregulatory domain. The RAVER1 protein contains an N-terminal domain with three RRMs and a C-terminal domain that functions as a splicing repressor. Our CTR is a fusion construct that contains N-terminal domain of *TARDBP* (encoding amino acid (AA) 1-268), with the NLS and both RRMs and the minimal repressor domain from *RAVER1* (encoding AA 433-626). The CTR also retains the 3’ autoregulatory domain from *TARDBP*. (**b**) The AAV-PHP.eB viral capsid, which can cross the blood-brain barrier, containing CTR transgene, is injected into control (*ChAT-Cre;Tardbp^f/+^*) or TDP-43cKO (*ChAT-Cre;Tardbp^f/f^*) mice through intravenous injection. (**c**) Schematic of the temporal analysis showing behavior and molecular analyses. (**d**) Breeding strategy for generating *ChAT-Cre;Tardbp^f/f^* (TDP-43cKO) mice which shows TDP-43 depletion in ChAT positive motor neurons and littermate control mice, *ChAT-Cre;Tardbp^f/+^*, which maintains the normal level of TDP-43. % indicates the Mendelian frequency of pups from each litter expected to carry the genotype of interest. (**e**) Breeding strategy for generating *ChAT-Cre;Tardbp^f/f^;Unc13aCE^-/-^*(TDP-43cKO with *Unc13aCE* KO) mice which shows TDP-43 depletion and *Unc13a CE* deletion in ChAT positive motor neurons and littermate control mice, which maintains the normal level of TDP-43. % indicates the Mendelian frequency of pups from each litter expected to carry the genotype of interest.

**Extended Data Fig. 2:**
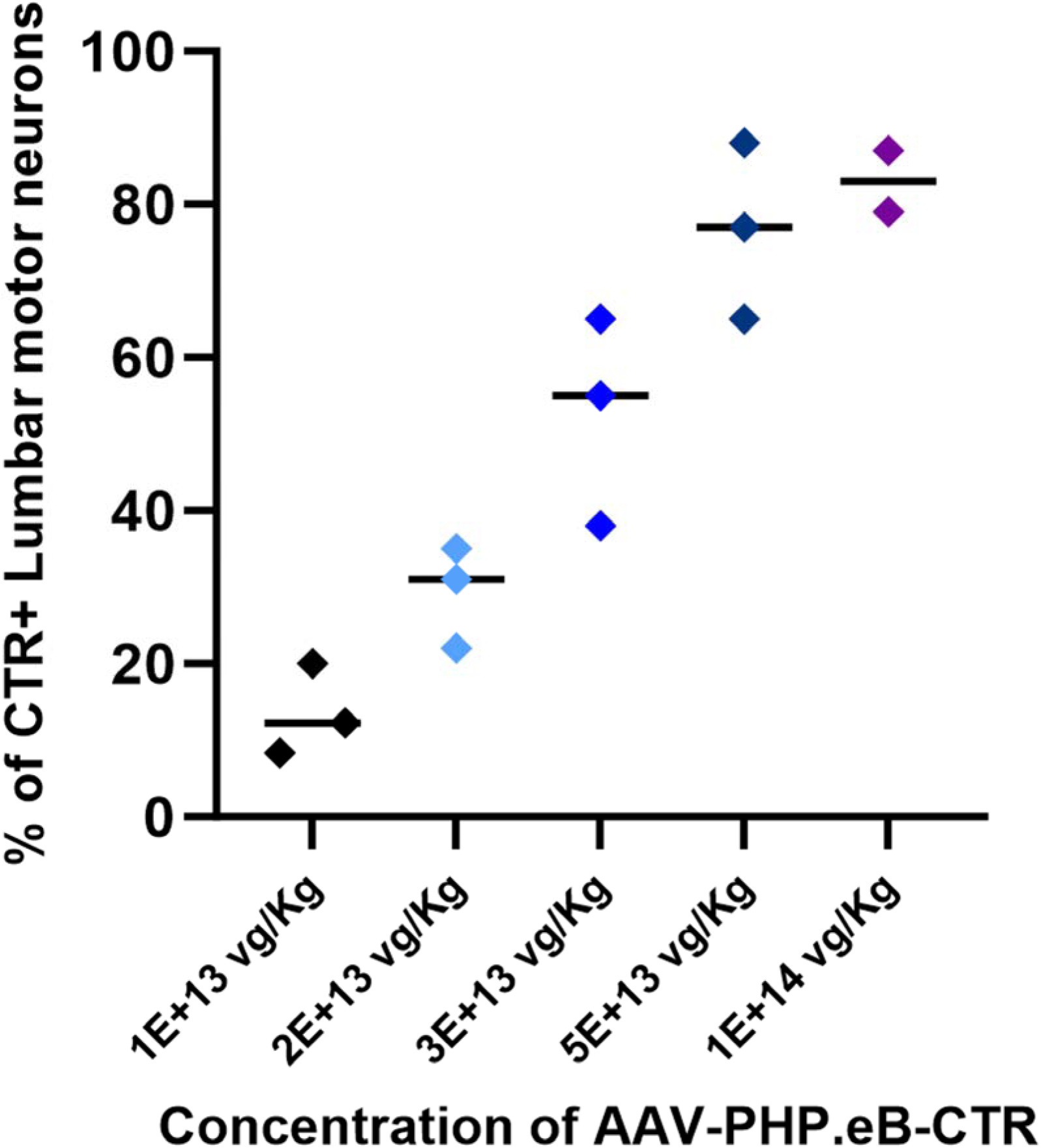
Optimization of the dosage of AAV-PHP.eB vector for intravenous delivery. The concentration of viral stock solution was 2E+13 vg/mL. Doses tested include 1E+13 vg/mL (n=3), 2E+13 vg/Kg (n=3), 3E+13 vg/Kg (n=3), 5E+13 vg/Kg (n=3), and 1E+14 vg/Kg (n=2). The optimum dose selected for the study was 5E+13 vg/Kg showing an average infectivity rate of 77%.

**Extended Data Fig. 3:**
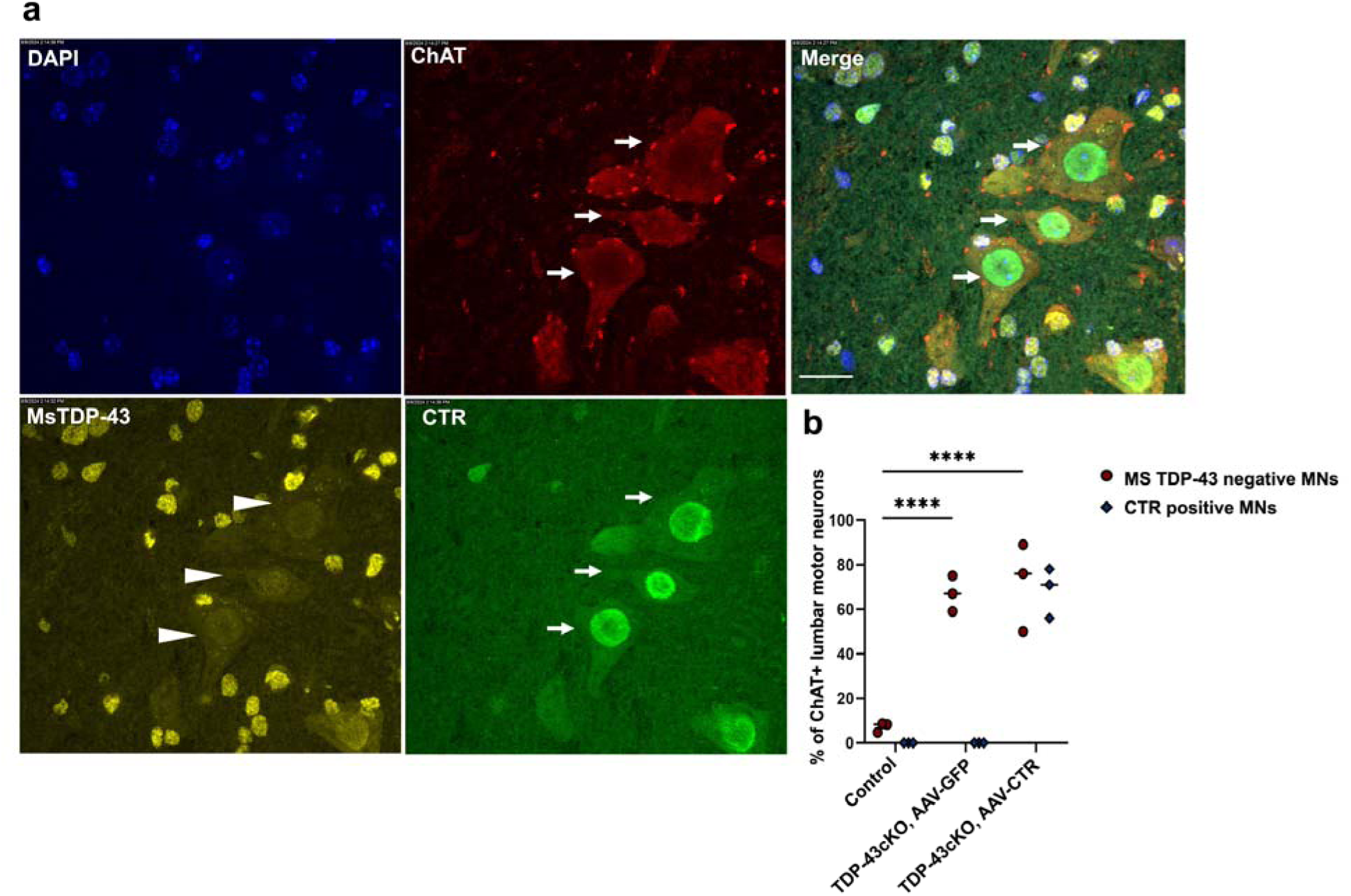
(**a**) Representative confocal images of lumbar spinal cord section from AAV-PHP.eB-CTR treated TDP-43cKO (*ChAT-Cre;Tardbp^f/f^*) mice, triple stained with human TDP-43 (CTR) (green), mouse endogenous TDP-43 (yellow) and ChAT (red). Nuclei were counterstained with DAPI (blue). Arrows indicate ChAT and CTR immunostaining in motor neurons and arrowheads show the depletion of endogenous TDP-43 in motor neurons. (**b**) Quantification of the percentage of lumbar motor neurons depleted of mouse TDP-43 and stained positive for CTR (n=3 per group).

**Extended Data Fig. 4:**
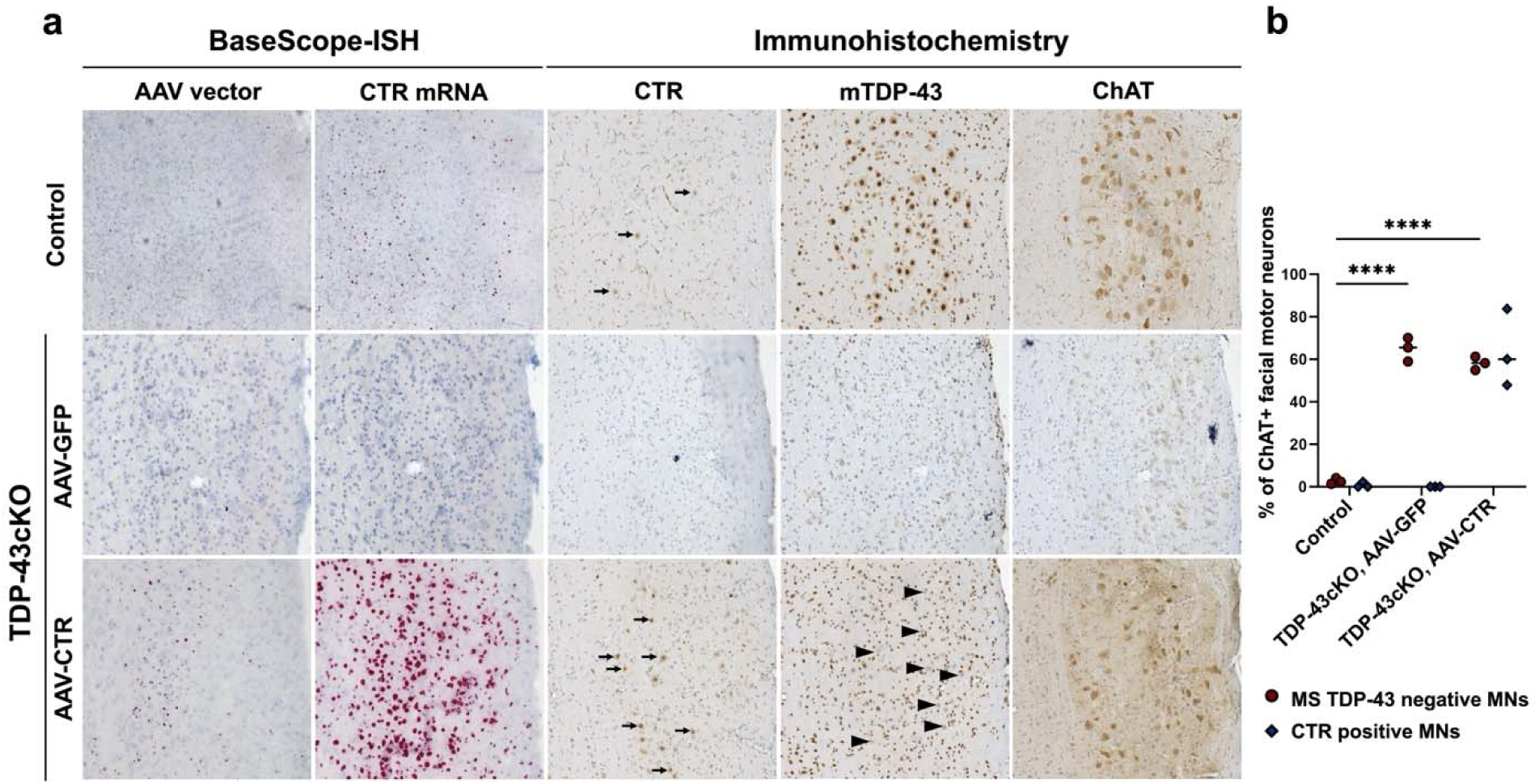
**(a)** Representative images showing accumulation of CTR in the facial motor nuclei of TDP-43cKO (*ChAT-Cre;Tardbp^f/f^*) mice. The intravenous administration of AAV-PHP.eB-CTR results in a mean infectivity rate of 72±11.7% (n=3). (**b**) Quantification of the percentage of motor neurons depleted of TDP-43 and stained positive for CTR. Arrows indicate anti-human TDP-43 immunostaining for CTR, and arrowheads indicate mouse endogenous TDP-43 depletion in TDP-43cKO mice. Panel 1. Control (*ChAT-Cre;Tardbp^f/+^*) mice, Panel 2. AAV-PHP.eB-GFP treated TDP-43cKO (*ChAT-Cre;Tardbp^f/f^*) mice and Panel 3. AAV-PHP.eB-CTR treated TDP-43cKO (*ChAT-Cre;Tardbp^f/f^*) mice. Scale bar=50 µm, n=3 per group.

**Extended Data Fig. 5:**
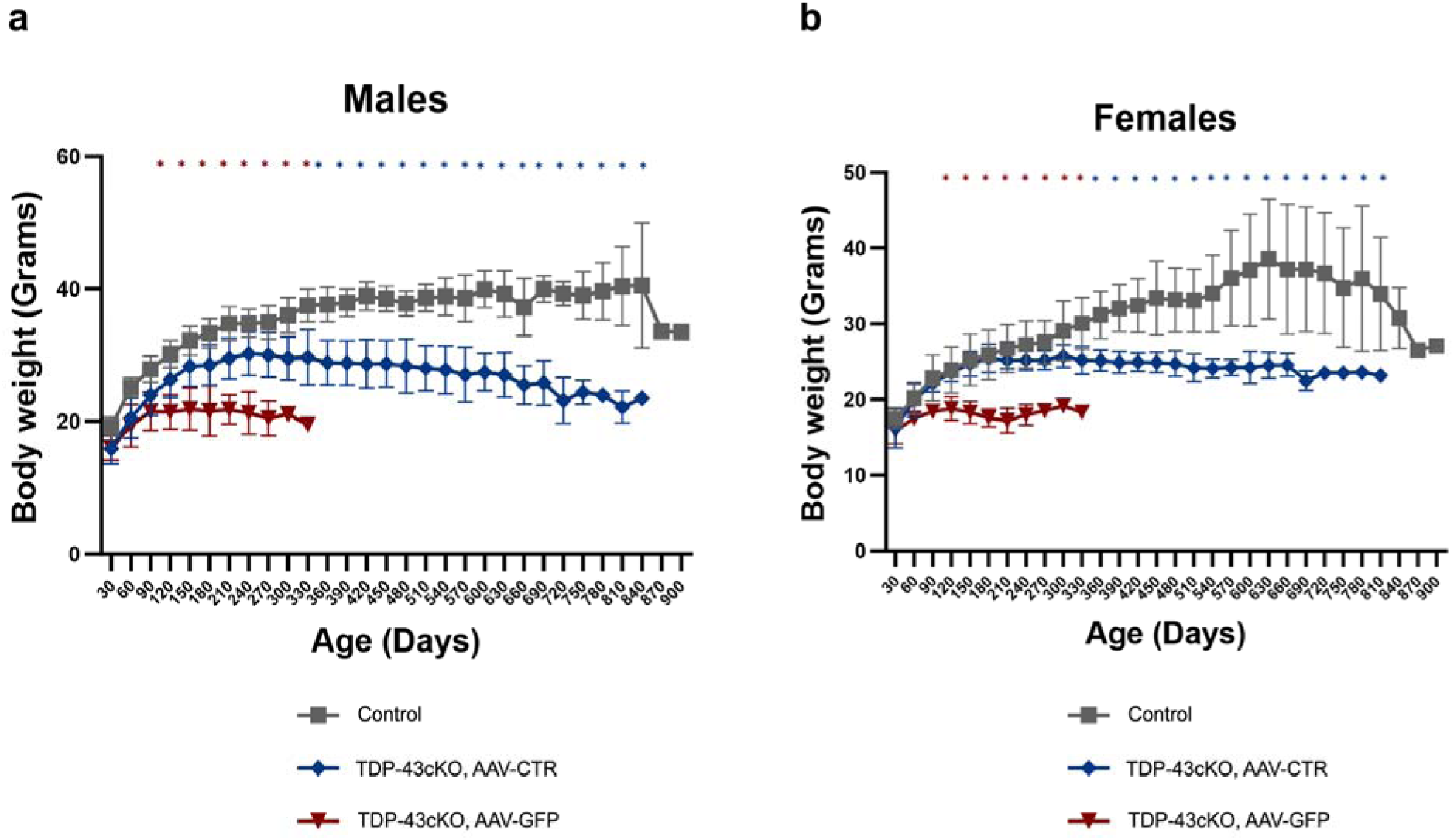
**Male and female body weight**

**Extended Data Fig. 6:**
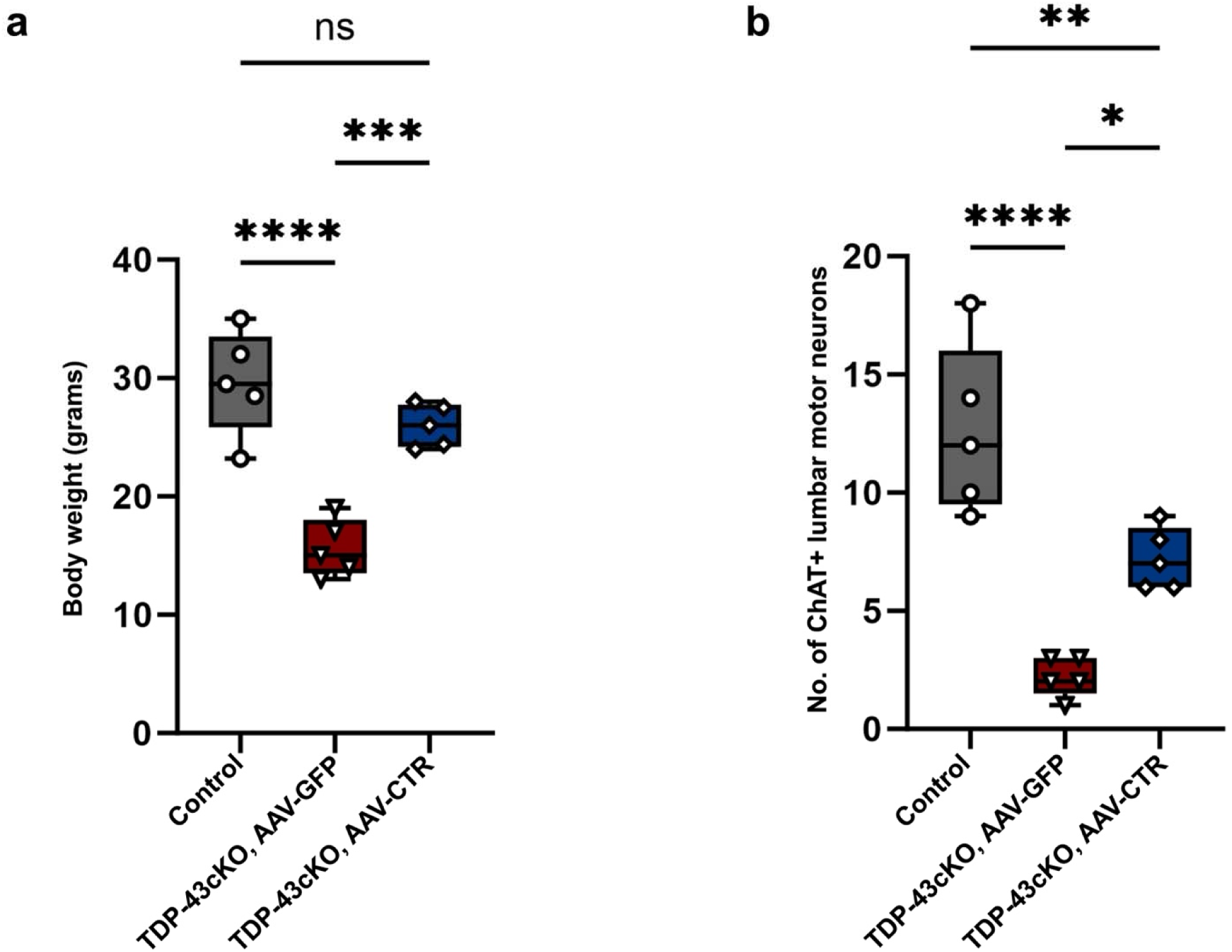
Body weight and lumbar motor neuron quantification in end stage mice (Control and TDP-43cKO mice treated with AAV.PHP.eB-CTR at 24-28 months of age and TDP-43cKO mice treated with AAV.PHP.eB-GFP mice at 8-12 months of age, n=5 per group).

**Extended Data Fig. 7:**
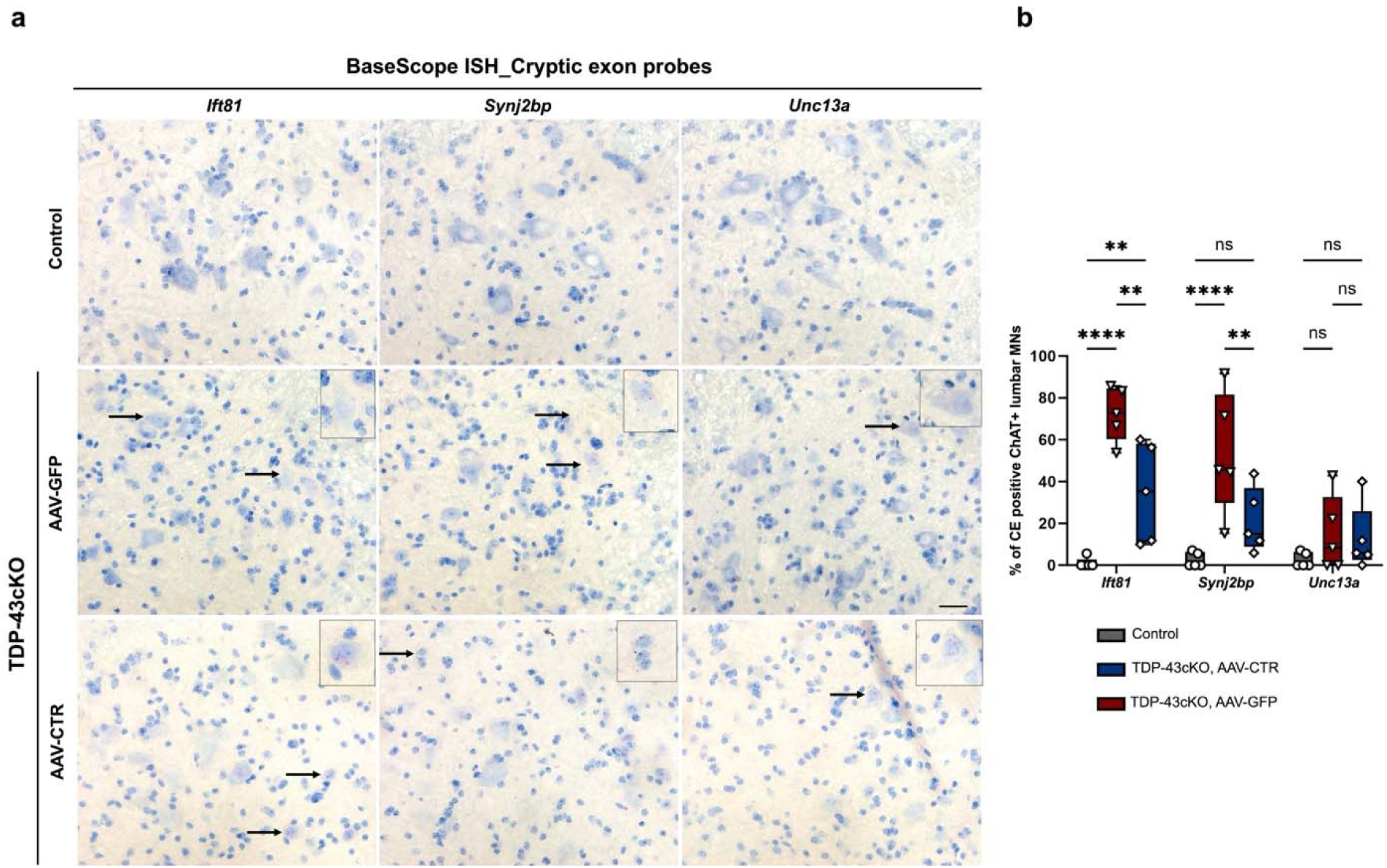
BaseScope ISH assay detecting TDP-43 associated cryptic exons in mouse spinal cord. (**a,b**) BaseScope ISH assay and quantification results using probes designed to detect cryptic exons in *Ift81*, *Synj2bp* and *Unc13a* demonstrates that the level of *Unc13a* cryptic mRNA was low in the TDP-43cKO mice treated with either AAV.PHP.eB-CTR or AAV.PHP.eB-GFP. Scale bar= 20 µm, n=5 per group.

**Extended Data Fig. 8:**
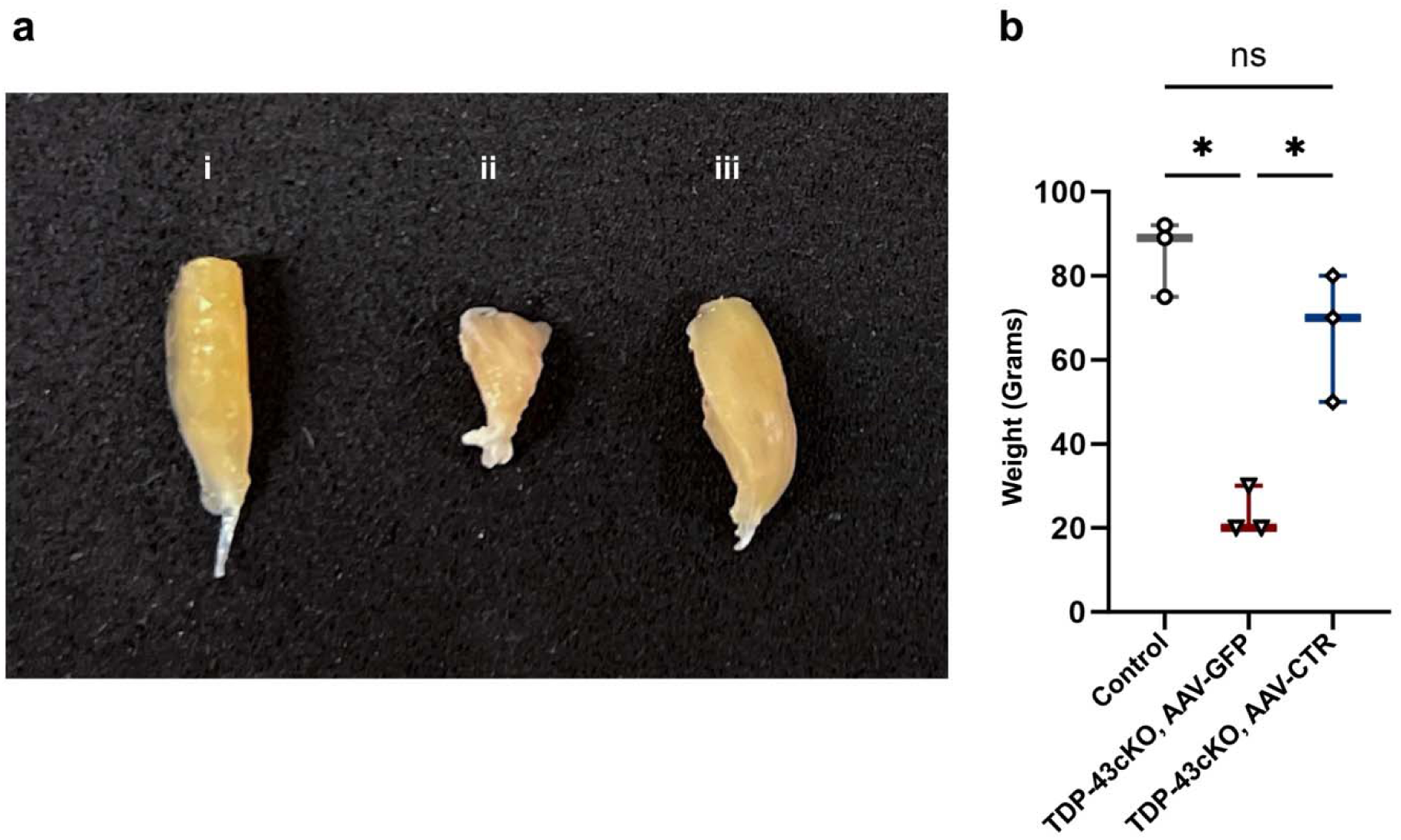
Gross analysis of hindlimb muscle size and weight shows rescue in TDP-43cKO mice treated with AAV-PHP.eB-CTR. (**a**) Representative image of the gross structure of gastrocnemius muscle in Control (i), TDP-43cKO mice treated with AAV-PHP.eB-GFP (ii) and TDP-43cKO mice treated with AAV-PHP.eB-CTR (iii). (**b**) Average weight of the gastrocnemius muscle was significantly reduced in TDP-43cKO mice treated with AAV-PHP.eB-GFP compared to the control mice (p value=0.006) and is rescued in TDP-43cKO mice treated with AAV-PHP.eB-CTR compared to TDP-43cKO treated with AAV-PHP.eB-GFP (p=0.006) (n=3 per group).

**Extended Data Fig. 9:**
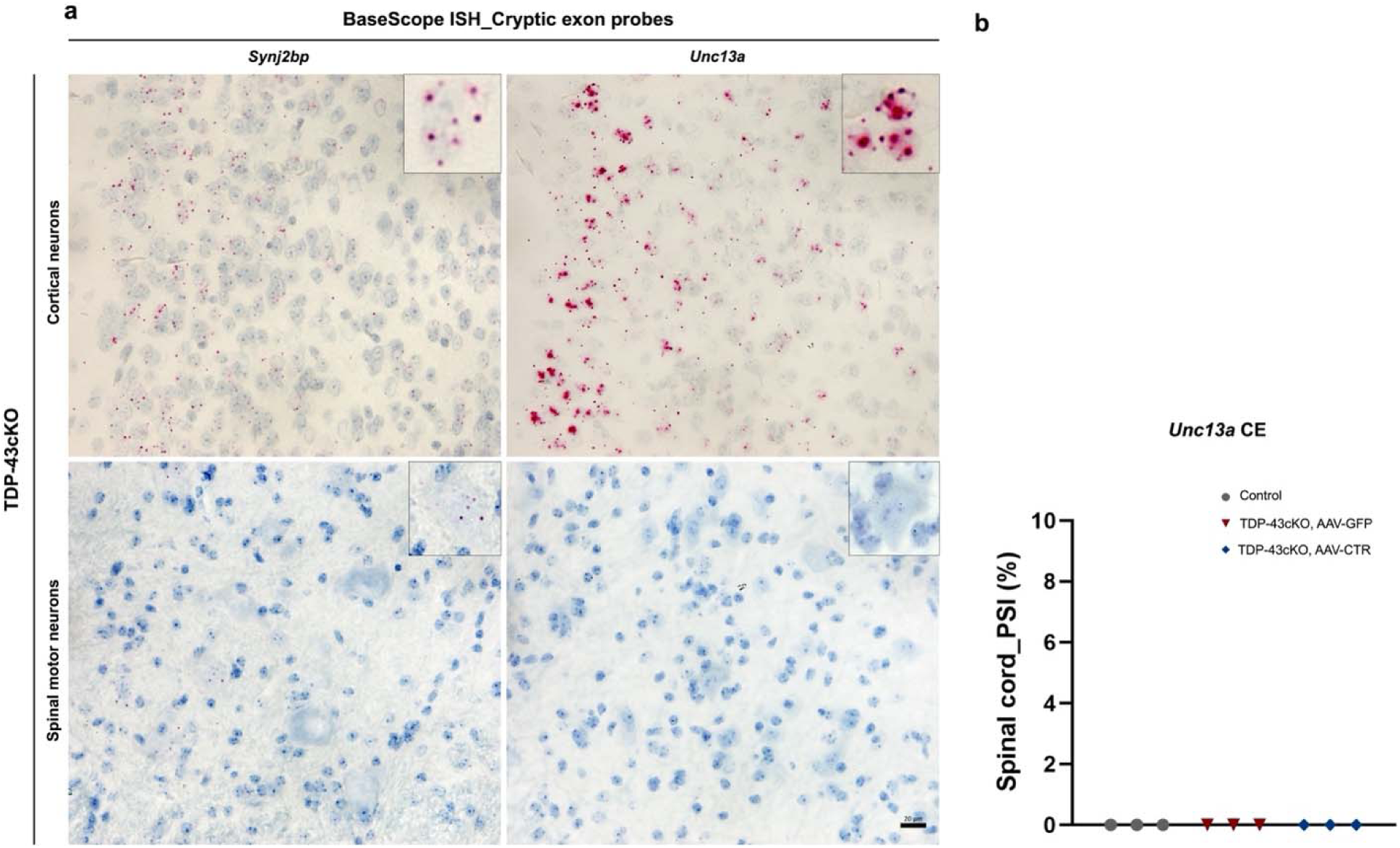
(**a**) Comparison of TDP-43cKO associated *Unc13a* and *Synj2bp* cryptic exon expression in the brain tissue (cortex, TDP-43cKO in excitatory neurons) and spinal cord (lumbar spinal cord, TDP-43cKO in motor neurons). TDP-43 depleted cortical neurons show robust staining for the *Unc13a* cryptic exons compared to the TDP-43 depleted spinal motor neurons. Scale bar= 20 µm. (**b**) PSI values from RNA-seq data shows that *Unc13a* cryptic exon expression is very low in spinal motor neurons (n=3 per group).

## ACKNOWLEDGMENT

We thank Michael Delannoy and Aswin Chandrasekhar for their technical assistance. This work was supported in part by the NIH grants (UG3/UH3NS115608, R33NS115161 and R01NS095969 to P.C.W.) the Kissick Family Foundation Frontotemporal Dementia Grant Program (to J.P.L.), and the Department of Defense (AL230138 to P.C.W.).

## AUTHOR CONTRIBUTIONS

P.C.W. and A.P.M. conceptualized, designed and interpreted the study. A.P.M., P.C.W. and J.P.L. wrote the manuscript. A.P.M., M.S.B., O.S., J.G.Y, T.C., S.R., I.R.S., T.M., and J.P.L., performed experiments and/or analyzed the data. All the authors reviewed and approved the final manuscript.

